# Predicting gene expression using millions of yeast promoters reveals *cis*-regulatory logic

**DOI:** 10.1101/2025.01.25.634853

**Authors:** Tirtharaj Dash, Susanne Bornelöv

## Abstract

**Motivation:** Gene regulation involves complex interactions between multiple transcription factors. While early attempts to train deep neural networks to predict gene expression were limited to naturally occurring promoter sequences, the advent of gigantic parallel reporter assays has expanded available training data by orders of magnitude. Despite these advances, a clear understanding of how to use deep learning to study gene regulation is still lacking.

**Method:** Here we investigate the complex association between gene promoters and expression in *S. cerevisiae* using Camformer, a residual convolutional neural network that ranked 4th in the Random Promoter DREAM Challenge 2022. We present the original model trained on 6.7 million random promoter sequences and investigate 270 alternative models to determine what factors contribute most to model performance. Finally, we use explainable AI to uncover regulatory signals.

**Results:** We show that Camformer accurately decodes the association between promoters and gene expression (*r*^2^ = 0.914 ± 0.003, *ρ* = 0.962 ± 0.002) and provides a substantial improvement over previous state of the art. Furthermore, we show that a much smaller model with approximately 90% fewer parameters than the original model can achieve a high predictive performance. Using Grad-CAM and in silico mutagenesis, we demonstrate that the model learns both individual motifs and their hierarchy. For example, while an IME1 motif on its own increases gene expression, the co-occurrence of a UME6 motif provides a switch to strongly reduce gene expression. Thus, deep learning models such as Camformer can provide detailed insights into *cis*-regulatory logic.

**Availability and Implementation:** The data and code used and developed in our experiments are publicly available at: https://github.com/Bornelov-lab/Camformer.

## 1 Introduction

Gene regulation is a fundamental aspect of life, in which cells control when and to what extent specific genes are activated or repressed. The gene regulatory process is tightly controlled and dynamically adjusted in response to internal and external signals. Many such signals reside in the gene promoter region, in the form of short sequence motifs that are recognised and bound by transcription factors (TFs). However, such TF motifs, or near-matching motifs, are frequent, and their effect on gene expression depends on the surrounding sequence context, including competition and cooperation between different TFs. Despite considerable efforts to map the gene-regulatory network through characterisation of individual TFs [Gerstein et al., 2012, Guo et al., 2012], fully decoding the *cis*-regulatory code remains an open challenge. One key limitation has been the lack of suitable data to study TF cooperativity and spacing rules, as the genome itself does not contain enough TF binding site pairs to draw statistically robust conclusions [Avsec et al., 2021b].

Recently, deep learning approaches, fuelled by the ever-increasing availability of large-scale genomic data, have emerged as a promising technique for studying gene regulation. DeepSEA [Zhou and Troyanskaya, 2015] pioneered the use of convolutional neural networks (CNNs) to identify regulatory single-nucleotide variants (SNVs) by predicting their effects on 166 chromatin-binding factors. DanQ [Quang and Xie, 2016] demonstrated that predictive performance was improved by integrating a bi-directional long short-term memory (LSTM), hypothesised to capture motif interdependencies. DeepAtt [Li et al., [2020] introduced category attention, which reduced model size while improving performance further. Beyond regulatory variants, Zrimec et al. 2020] trained a model on 20,000 mRNA datasets, revealing how mRNA and flanking DNA sequences interact and associate with gene expression. BPNet [Avsec et al., 2021b] employed a CNN to provide base-resolution insights into binding and cooperativity of four key TFs. Finally, Enformer [Avsec et al., 2021a] used a combination of convolutional blocks and transformer blocks to handle longer sequences, enabling it to predict long-range regulatory interactions between enhancers and promoters.

However, these models were all trained on naturally occurring genomic sequences, which imposes a strong upper limit on the size and diversity of available training data. Massively parallel reporter assays (MPRAs) offer a potential solution, by enabling sequencing-based readouts of the activities of thousands of candidate regulatory elements in a single experiment [Patwardhan et al., 2009]. Traditionally, such assays used a pre-defined set of sequences. By instead using random sequences, gigantic parallel reporter assays (GPRA) have effectively enabled a potentially unlimited source of training data [de Boer et al., 2020, Vaishnav et al., 2022]. Conducting a GPRA, de Boer et al. [2020] constructed a library of over 100 million (M) random promoters and measured their ability to drive the expression of yellow fluorescent protein (YFP) in yeast (*S. cerevisiae*). This study, along with a follow-up study [Vaishnav et al., 2022], constitutes a valuable source of promoter-to-expression benchmarking data, facilitating the development and comparison of deep learning methods for studying the *cis*-regulatory code.

Community-wide challenges have been shown to drive the development of computational methods through systematic benchmarking. For instance, AlphaFold [Jumper et al., 2021] popularised and transformed protein structure prediction by winning the Critical Assessment of Structure Prediction (CASP) competition in 2018. Similarly, the Dialogue for Reverse Engineering Assessment and Methods (DREAM) competition aims to improve our understanding of biological processes through advancement in bioinformatic methods [Meyer and Saez-Rodriguez, 2021]. The Random Promoter DREAM challenge held in 2022 (DREAM2022) specifically sought to advance deep learning methods for predicting gene expression from promoter sequences. The participants were provided with GPRA data representing 6,739,258 promoters and their YFP expression levels. The goal of the challenge was to develop a model able to predict the expression of another 71,103 promoters, whose expression had been measured separately and were withheld from the participants during the competition. In total, 19 of the submitted solutions outperformed a benchmark transformer model, which represented previous state of the art [Vaishnav et al., 2022, Rafi et al., 2024]. Among these top models were LegNet [Penzar et al., 2023] in 1st place, Proformer [Kwak et al., 2024] in 3rd place, and CRMnet [Ding et al., 2023] in 12th place.

Camformer secured 4th place in DREAM2022 by designing a CNN with residual connections. Here, we provide an in-depth analysis of the submitted Camformer model, including a systematic grid search to pinpoint factors influencing model performance. Additionally, using model explainability techniques, we analysed the 71,103 sequences in the challenge test set, revealing both known and potentially novel gene-regulatory motifs. Overall, our study demonstrates that a carefully designed CNN can achieve state-of-the-art performance on gene-regulatory tasks, while providing important insights into the *cis*-regulatory code.

## 2 Materials and methods

### 2.1 Training data and evaluation

For completeness, we here provide a brief description of the DREAM2022 data. A more elaborate description of the experimental data can be found in the DREAM2022 paper [Rafi et al., 2024]. In short, Rafi et al. [2024] conducted a high-throughput GPRA to determine the regulatory effect of millions of promoters on gene expression. A library of 80-nucleotide (nt) random DNA sequences was cloned into a minimal promoter construct located upstream of a YFP and followed by a constitutively expressed red fluorescent protein (RFP). The resulting plasmid was transformed into yeast, its promoter activity was measured as log(YFP*/*RFP) by flow cytometry, followed by sorting the cells into 18 expression bins that were sequenced separately. An expression value was assigned to each promoter, describing its weighted mean expression bin.

In total, the public training set consisted of 6, 739, 258 promoter sequences along with their expression (mean of ∼2 cells per promoter) (Figure 1a,b). A test set with 71, 103 promoter sequences was sequenced separately, at higher depth, enabling a more precise estimation of their expression (mean of ≥100 cells per promoter). To enable a detailed evaluation of the submitted models, the test set included promoter sequences with different properties: sequences with SNVs (*n* = 44, 340 pairs), perturbation of known TF binding sites (*n* = 3, 287 pairs), tiling of known TF binding sites across random sequences (*n* = 2, 624 pairs), sequences with high (*n* = 968) and low (*n* = 997) expression levels, native yeast promoter sequences (*n* = 997), challenging sequences (*n* = 1, 953), and random sequences (*n* = 6, 349). The test set was kept private during the competition, with the exception of 13% that was used for the public leaderboard, where only *r*^2^, *ρ*, and weighted Pearson and Spearman scores were shown, keeping the performance on the different subsets hidden.

**Figure 1:**
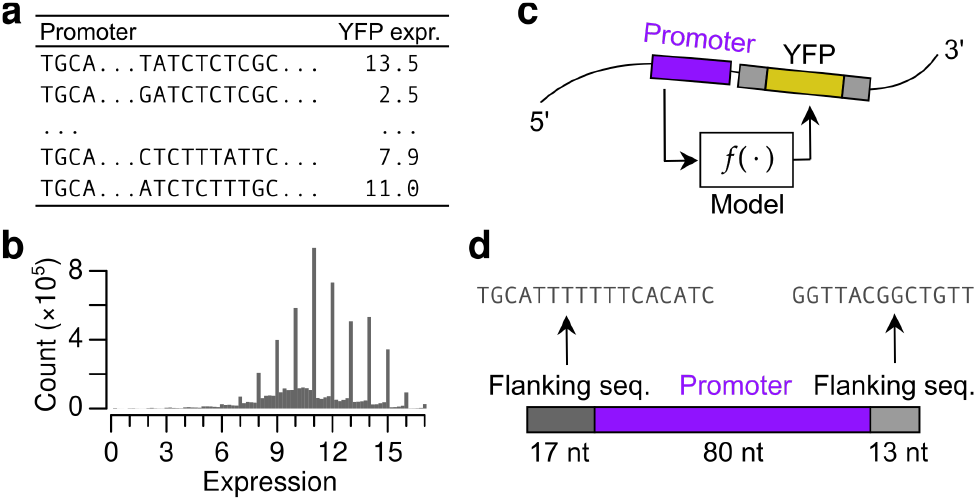
A schematic representing the prediction task and data: (a) The training data is arranged as (promoter, expression) pairs; (b) Histogram of gene expression in training set; (c) The task is to predict the expression of YFP given a promoter sequence; (d) Representing the promoters using a 110 nt fixed-length representation, the original 80 nt promoter is squeezed between two fixed flanking sequences of lengths 17 and 13 nts.

### 2.2 Prediction task

The promoter-to-expression prediction task can be stated as in the following definition. In our study, we explore how to define *f* using a CNN (Figure 1c).

#### Definition 1

*Let S* = {*A, C, G, T, N*} ^110^ *denote a promoter sequence of length* 110, *Here, A, C, G, T are the four nucleotides and N represents an unknown nucleotide. The gene expression prediction task is then to learn a mapping f* : *S* → ℝ.

### 2.3 Data preprocessing

Each promoter in DREAM2022 was represented by its ∼80 nt variable region inserted between two fixed sequences of length 17 and 13 nts respectively, representing the flanking plasmid sequence (Figure 1d). For training, we retained only sequences of length 110 ± 3 nts (discarding 110,138 sequences) and containing at most three *N* (discarding 23,533 sequences). Sequences shorter than 110 nts were padded with *N* at the end, and sequences longer than 110 nts were truncated.

## 3 Algorithm and implementation

Although the trending approach in sequence learning is using transformers [Lin et al., 2022], there have been numerous successful applications of CNNs in genomics [Avsec et al., 2021a, Zhang et al., 2021]. Here, we seek to demonstrate that a carefully designed CNN is sufficient to achieve state-of-the-art performance in predicting gene expression from promoter sequences. We base our reasoning on the fact that CNNs have an inherent ability to extract sequence motifs from input DNA sequences. If each kernel is seen as a position weight matrix (PWM), then the computations at a convolutional layer corresponds to calculating the PWM score across the input sequence, highlighting influence of each position in the input on the model’s prediction. This is akin to what transformers are explicitly made to do with the help of the attention mechanism [Vaswani et al., 2017].

### 3.1 Submission model

We first sought to determine an optimal structure for the CNN, which is often difficult. There are two ways to do this: (a) a classical grid search over different configurations or (b) using an automated Bayesian search. As a Bayesian search allows exploration of a larger search space, this approach was used for the competition. Nevertheless, to gain a better insight into model design choices, we later performed systematic experiments using a grid search, and elaborate on this in the subsequent sections.

### 3.1 Sequence encoding

Preprocessed training sequences (*n* = 6, 605, 587) were one-hot encoded using the following encodings: *A* = [1, 0, 0, 0], *C* = [0, 1, 0, 0], *G* = [0, 0, 1, 0], *T* = [0, 0, 0, 1], and the special symbol *N* = [0, 0, 0, 0]. Test data (*n* = 71, 103 sequences) were one-hot encoded in the same manner. Hence, each promoter sequence is uniformly represented as a tensor of dimension (*height*[*h*], *width*[*w*], *channels*[*c*]) = (110, 1, 4).

#### 3.1.2 Training loss function

Since the training data were expected to be noisy, we used mean absolute error (MAE), also called *L*_1_ loss, between true and predicted expression as the loss function during training. This allows the outliers to be handled in a more robust manner.

#### 3.1.3 Optimiser

We used the Adam optimiser with weight decay regularisation (AdamW). Maximum number of epochs was set to 50, but in practice, we used early-stopping to obtain a well-generalised model by monitoring performance (*r* + *ρ*) on a validation set (8 % of the training data).

#### 3.1.4 Model refinement

Construction of candidate models is described in Supplementary Methods 1.1.1. Candidate models were refined using a Bayesian search through the Optuna hyperparameter optimisation software [Akiba et al., 2019], specifically designed for automated machine learning. Optuna was used to vary batch size, learning rate, weight decay, size of the convolutional kernels, number of convolutional and fully connected layers, number of channels and dropout rates (Figure 2a). Most runs were performed on subsets of either 1 or 2 M promoters, and allowing up to 100 models to be tested. Promising models were then re-trained on the full training set. We noticed that several independent runs converged towards adding a single max pooling operation at the penultimate layer. Indeed, although our Optuna run that returned our ResNet-inspired model did not explore any pooling operations, adding it manually improved model performance.

**Figure 2:**
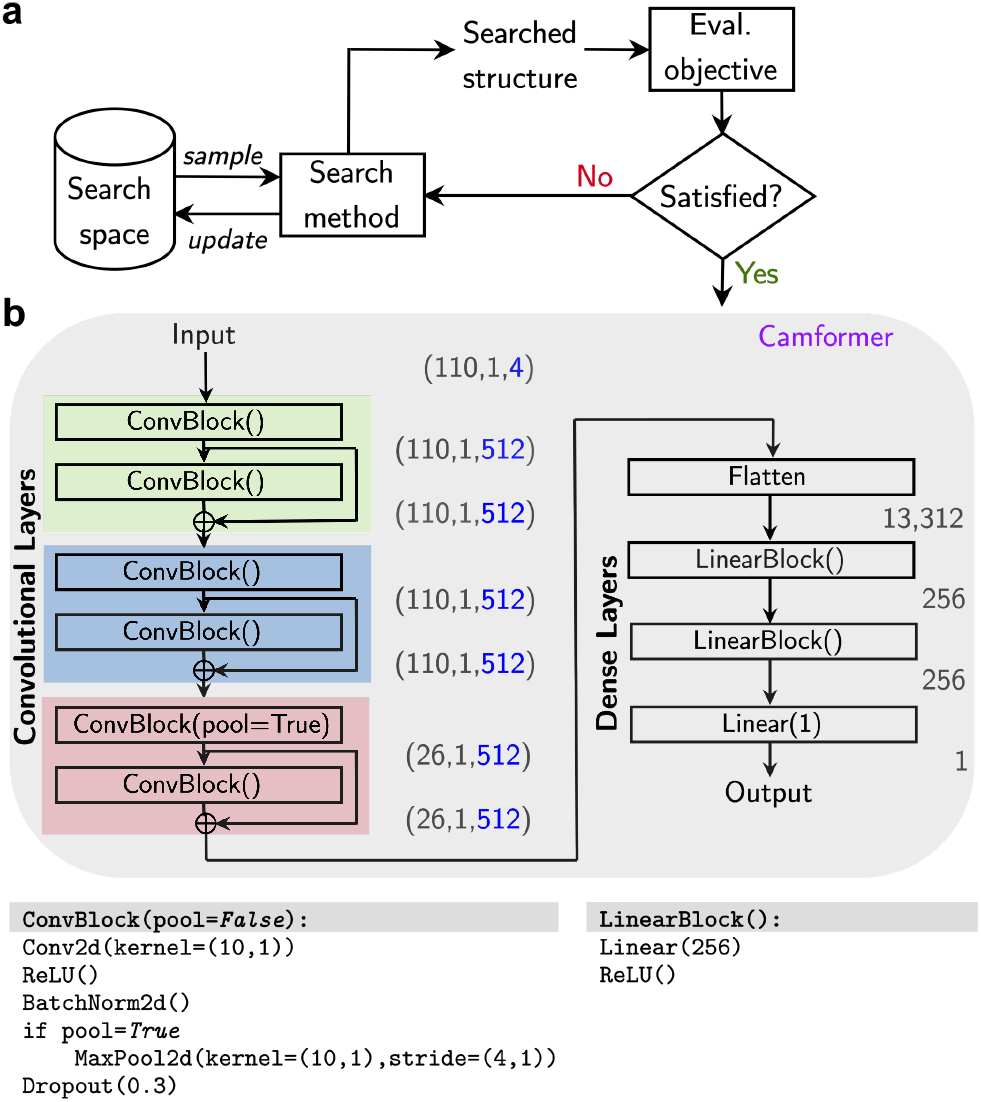
Overview of the search for an optimal model architecture. (a) We used Optuna [Akiba et al., 2019] to perform a hyperparameter search over a space of CNN architectures; (b) The final model consists of six convolutional layers followed by fully connected layers. Inputs and activations are shown in the format (*h, w, c*).

#### 3.1.5 Camformer

The model submitted for DREAM2022 is shown in Figure 2b and referred to as ‘Camformer’. The final model has six convolutional layers with residual connections after layers 2, 4, and 6, followed by three fully connected layers to compute the predicted expression. Overall, the model had 16.6M trainable parameters.

### 3.2 Hyperparameter grid search

To determine what factors contributed to model performance, we performed a systematic grid search over model structures, different types of sequence encoding strategies, loss functions, optimisers, and learning rate schedulers. Due to the systematic evaluation performed, this experiment allowed us to probe the effect of each of the tested variables. To keep the experiment tractable, we restricted ourselves to three basic structures of Camformer: Camformer, Camformer-Small and Camformer-Mini. These models and the full grid search is described in Supplementary Methods 1.2.

### 3.3 Evaluation

#### 3.3.1 Quantitative, predictive evaluation

We used a subset of the training set and report their empirical performance on a test set, which is a smaller subset of the training set. We adopt the Pearson and Spearman correlation coefficients (*r* and *ρ*, respectively) between true and predicted expressions for evaluating the models. The former is more sensitive to outliers than the latter. Evaluations were carried out against all nine categories of promoter sequences in the test set (refer to Section 2.1). In addition, we also calculate the *Pearson Score* and *Spearman Score* metrics used in the competition to rank different submissions, which is a weighted average of performance across different categories of promoters in the test set. Overall, 20 different evaluation metrics are computed for each model.

#### 3.3.2 Qualitative, explanatory evaluation

Here we assess Camformer in the context of model explainability, aiming to identify which promoter features the model is attending to when making its predictions. For this, we use the following two approaches (more details in Supplementary Methods 1.3):

1. Gradient-weighted class activation maps (Grad-CAM): Grad-CAM [Selvaraju et al., 2017] highlights important regions in the input sequence by analysing the gradients of the model’s output with respect to the input to each convolutional layer.
2. In silico mutagenesis (ISM): ISM measures how the model responds to variations in the input sequence by systematically mutating each position of the input sequence and observing the change in the predicted output.

We further visualise the information generated by these two approaches using sequence logos, with an intention to highlight important TF motifs in the promoters. While both approaches might highlight similar motifs, the logos from the ISM could offer a more comprehensive view of the sequence positions critical to the model, while Grad-CAM might emphasise the most salient features.

### 3.4 Hardware

During the competition, all models were developed on the Google TPU Research Cloud (TRC) on TPU v2-8 or v3-8. We used either PyTorch (tpu-vm-pt-1.11) or TensorFlow (tpu-vm-tf-2.8.0). Subsequent work was performed on a Dell workstation with Ubuntu 22.04 LTS, Intel i9-13900K processor, 64GB memory, NVIDIA GeForce RTX 4090, using PyTorch 2.1.1 with NVIDIA CUDA toolkit version 12.1 and Python 3.11. More details on other relevant libraries used in our work can be found in the code repository.

## 4 Results

### 4.1 Camformer robustly predicts gene expression

Camformer placed 4^th^ out of 110+ teams in the DREAM2022 challenge [Rafi et al., 2024]. Notably, its reported performance (*r*^2^ = 0.913, *ρ* = 0.961) provided a substantial improvement over the reference model [Vaishnav et al., 2022] at 20^th^ place (*r*^2^ = 0.879, *ρ* = 0.938). However, this evaluation did not assess the stability of model design and training strategy. We therefore re-trained the Camformer model 10 times using different random seeds. Out of the 10 models, more than half of them exceeded previously reported performance (*r*^2^ = 0.914 ± 0.003, *ρ* = 0.962 ± 0.002, Figure 3a-c, Figure S2), showing that the model is not only accurate but also robust.

**Figure 3:**
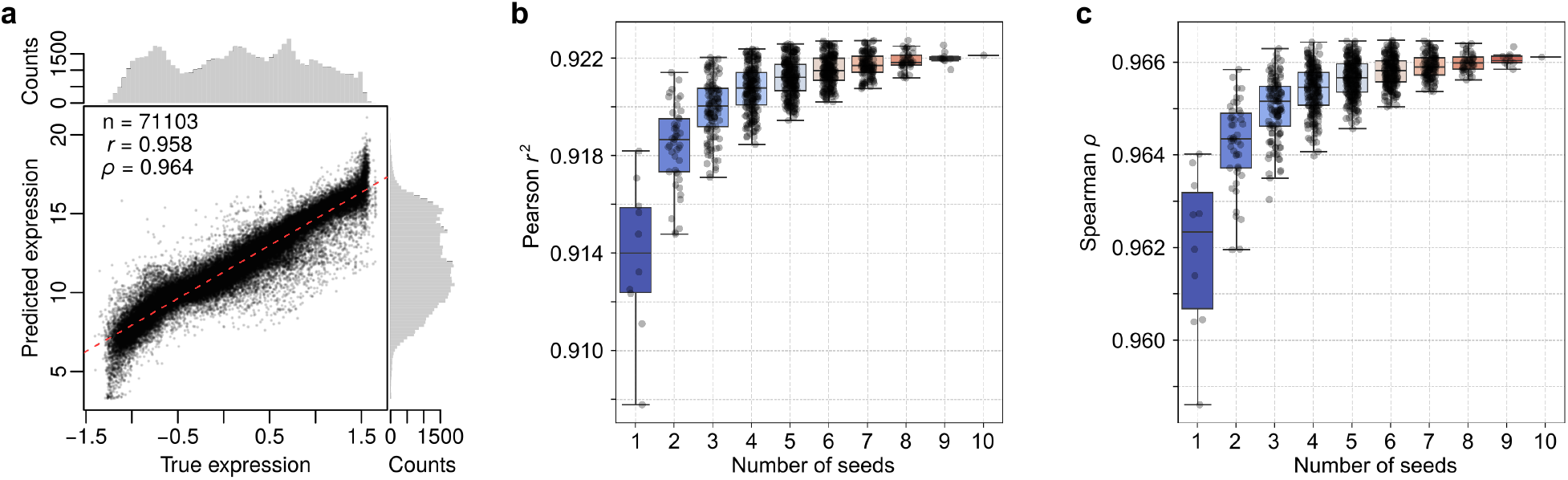
Camformer accurately predicts gene expression from promoter sequence. The model was evaluated on the DREAM2022 test dataset (*n* = 71, 103). (a) Scatter plot between true and predicted expression values, (b) Predictive performance measured using Pearson *r*^2^ across ten individual replicates and ensemble models constructed by taking the mean prediction from *k* replicates (where *k* ∈ {2, …, 10}). A replicate here refers to a model constructed using a random seed, determining the initialisation of model parameters (weights) before training. (c) Same as b), but showing Spearman *ρ*. Please note that the range of expression values in the test dataset differed from that of the training dataset.

To investigate how model design influences its performance, we repeated the same experiment for Camformer-Mini, a small CNN (1.4M parameters) without residual connections, which was developed during the competition and later optimised (described below in Section 4.1.1). Although Camformer-Mini displayed lower performance (*r*^2^ = 0.898 ± 0.004, *ρ* = 0.952 ± 0.003), it nevertheless exceeded previous state of the art (Figure S3a-c).

A common strategy to improve prediction performance is to use an ensemble model, combining predictions from multiple models. Although this strategy was not allowed in the competition, both Camformer and Camformer-Mini displayed significant performance gain and reduced variance when an ensemble was constructed by averaging predictions from two or more seeds (Figure 3b-c, Figure S3b-c). For instance, using two seeds significantly improved Camformer *r*^2^ from 0.914 ± 0.003 to 0.918 ± 0.002 (*p*=0.001, two-sided Student’s t-test) and *ρ* from 0.962 ± 0.002 to 0.964 ± 0.001 (*p*=0.002). As expected, the best performance was observed using all ten seeds (*r*^2^ = 0.922, *ρ* = 0.966). Thus, even without changing model design, performance can readily be improved by re-training the same model several times with different parameter initialisations.

#### 4.1.1 Optimiser and sequence encoding contribute to model performance

To determine what factors contribute to model performance, we performed a hyperparameter grid search. For this experiment, we considered Camformer (16.6M parameters) and Camformer-Mini (1.4M), and additionally a small variant of the Camformer, Camformer-Small (3.4M), where the number of channels was reduced from 512 to 256 at each convolutional layer. In addition to model structure, we explored sequence encoding, loss function, optimiser, and LR scheduler (see Supplementary Methods 1.2.1). In total, 90 hyperparameter combinations were evaluated for each model structure, resulting in 270 models.

Notably, the AdamW optimiser improved median *r* by 4.3% compared to Lion (*p*=7e-24, two-sided Wilcoxon Rank Sum test, n=145 model comparisons) as shown in Figure S4a. We therefore analysed the two optimisers separately (Figure S4b,c). Using AdamW, both Camformer and Camformer-Small performed significantly better than Camformer-Mini (median *r* increased by 0.4-0.5%, *p <* 5e-6, *n* = 45). Moreover, across the five sequence encodings tested, encodings without an uncertainty channel (‘onehot’, ‘onehotWithP’ and ‘onehotWithN’) consistently performed better than those with uncertainty (‘onehotWithInt’ and ‘onehotWithBoth’), displaying a median *r* increase by 0.5-0.6% across all comparisons (*p <* 2e-5, *n* = 27). The same trend was seen for Lion, although the gain was more variable (0.5-2%) and not always significant. Full results for the grid search are available on our GitHub repository.

We next selected the best set of hyperparameters for each model structure and compared the resulting model to the original Camformer model. For this, we re-trained each selected model ten times using the whole training set and evaluated it on the external test set. Evaluating the models across nine categories of promoter sequences demonstrated that the original Camformer model remained the best model (Figure S5-S7). We therefore used Camformer for the remaining analyses, unless otherwise specified.

#### 4.1.2 Challenging sequences and motif perturbations display strongest improvements

To better understand its performance, we compared Camformer to the benchmark transformer model by Vaishnav et al. [2022]. However, benchmark performance differed substantially across different sequence categories, ranging from *r*^2^ = 0.264, *ρ* = 0.305 for high expressed sequences to *r*^2^ = 0.912, *ρ* = 0.956 for random sequences (Figure S5, Figure S6). Since neither *r*^2^ nor *ρ* can exceed 1, the maximum possible improvement therefore ranges from 278% and 227% for high expressed to 9.7% and 4.6% for random sequences, which would both result in an *r*^2^ and *ρ* of 1.

To provide a fair comparison of the improvement against different baselines, we therefore used baseline-normalised improvement, defined as Δ^*norm*^*ρ* = Δ*ρ/*(1 − *ρ*_1_) = (*ρ*_2_ − *ρ*_1_)*/*(1 − *ρ*_1_). Normalised improvement in *ρ* was 36.9% for all sequences, and ranged from 8.2% for SNVs, 13.5% for native, 29.5% for motif tiling, 30.3% for high expressed, 31.9% for low expressed, 36.1% for random sequences, 39.5% for motif perturbation and 56.8% for challenging sequences (Figure S5). While normalised performance across most categories was increased around 30-40%, the most notable exceptions were the strong improvement (56.8%) on challenging sequences (Figure 4a) and the weak improvement (8.2%) on SNVs (Figure 4b). Notably, the LegNet model also provided its lowest improvement at 19.3% for SNVs, potentially reflecting an inherent difficulty for a model trained to predict the expression of diverse random sequences to understand the meaning of SNVs (Figure 4b).

**Figure 4:**
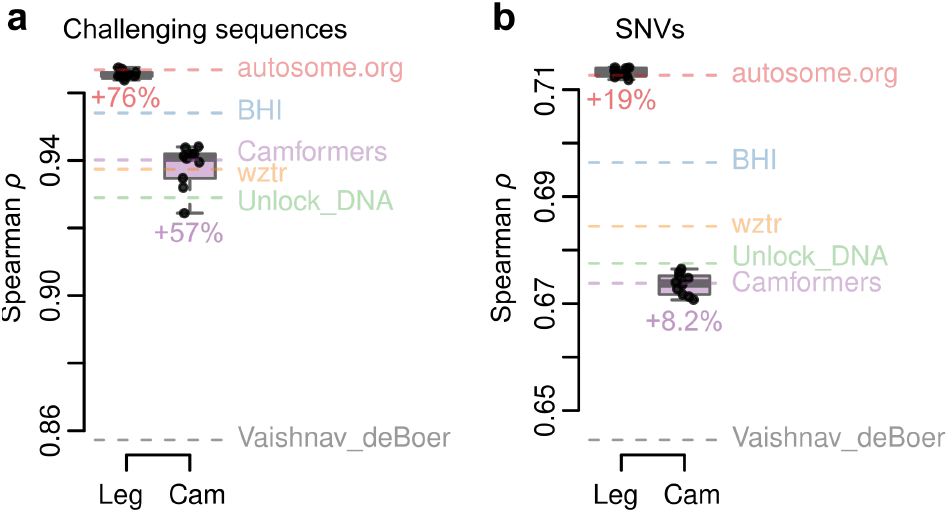
Performance of Camformer (Cam) and LegNet (Leg) across (a) challenging sequences, which displayed the strongest improvement (arrows) over baseline (Vaishnav_deBoer; dashed grey line), or (b) SNVs, which displayed the weakest improvement over baseline. The top five DREAM2022 submissions are shown as coloured dashed lines. Results for all test set categories are shown in Figure S5.

#### 4.1.3 Predictive performance near state-of-the-art

Vaishnav et al. [2022] previously released two additional datasets, representing yeast grown in complex (YPD) or defined (SD-Ura) media. Each dataset contains promoter sequences and their YFP expression measured using the same type of GPRA assay as the one used to train Camformer. The complex media data included 30,722,376 training and 3,929 test sequences, whereas the defined media had 20,616,659 training and 3,978 test sequences. In each case, the training data consisted of random sequences and the test sequences represented fragments from native yeast promoters, both 80 nts long and flanked by the same 17 and 13 nts as before, making the total length 110 nts. Despite these data representing yeast grown under different conditions, we first tested how our existing Camformer and Camformer-Mini models performed on the two test sets. Notably, Camformer achieved high performance (*r* = 0.956, *ρ* = 0.964 for complex media; *r* = 0.950, *ρ* = 0.958 for defined media) (Figure 5b,d), similar to LegNet (*r* = 0.960 and *r* = 0.950 for complex and defined media, respectively).

**Figure 5:**
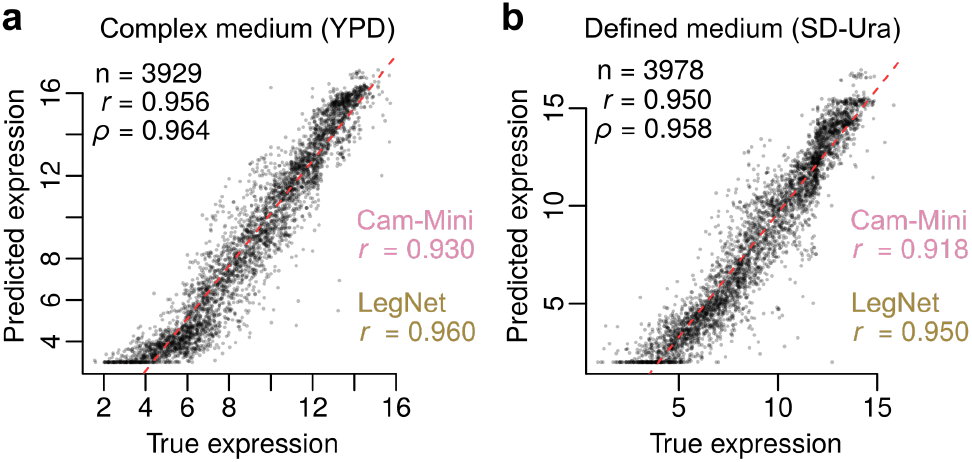
Performance of Camformer in predicting expression from promoters in yeast grown in (a) complex medium (YPD) and (b) defined medium (SD-Ura). Performance of Camformer-Mini and LegNet [Penzar et al., 2023] is provided as a comparison (pink and yellow).

We next re-trained Camformer and Camformer-Mini using the training data from the complex and defined media. To understand the relationship between training data size and model performance, we performed a sub-sampling experiment, using increasingly larger training data with 2M increments. We noticed a steady improvement with increasing training data size for Camformer, in contrast, Camformer-Mini displayed no clear improvement with additional training data, suggesting that Camformer-Mini may not be sufficiently parameterised to benefit from more training data (Figure S8a,b).

Using Camformer and Camformer-Mini trained on complex media, we compared their performance to currently leading models, including DeepSEA, DanQ, DeepAtt, the benchmark model developed by Vaishnav et al. [2022], and LegNet, the DREAM2022 winner, all re-trained using the same data (Figure 6a,b). Notably, Camformer provided substantial improvement over DeepSEA, DanQ, and DeepAtt, as well as over the benchmark model, and was only marginally worse than LegNet for native yeast sequences (*r* = 0.956), while having a strong performance just above the benchmark also on random sequences (*r* = 0.968). We therefore conclude that the Camformer archtecture provides state-of-the-art performance also on very large training sets.

**Figure 6:**
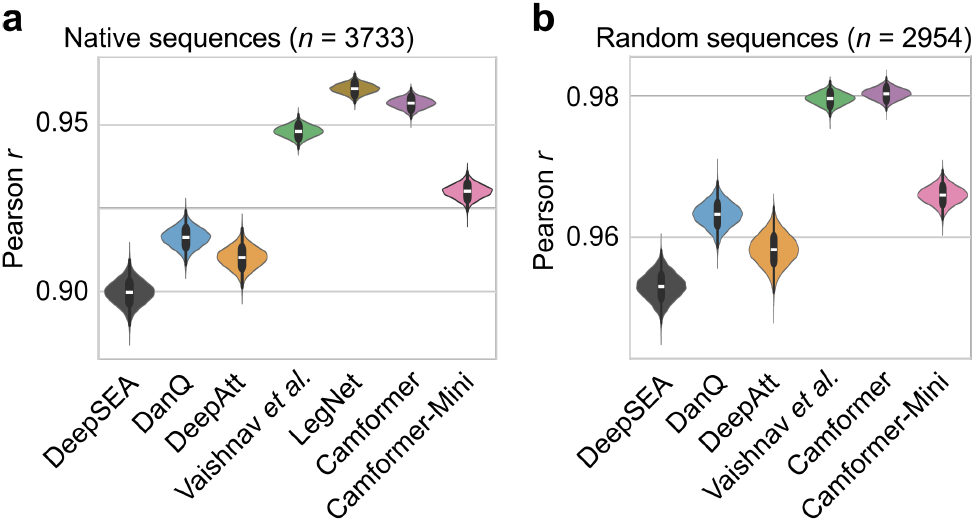
Comparison of Camformer and Camformer-Mini with the state-of-the-art models. Prediction of native promoter expression for yeast grown in complex medium (YPD) as shown in (a) and random promoter expression, as shown in (b). Violin plots show bootstraps with *n* = 10, 000 replicates. All model differences were statistically significant (*p <* 0.05, paired Student’s t-test).

### 4.2 Sequence embeddings reveal decision process

We next studied the internal sequence embeddings learnt by Camformer. We randomly sampled 10,000 sequences from the test set and passed them on to the model. For each convolutional layer, the outputs are tensors of dimension (*h*, × *w*, × *c*). To visualise these sequences using t-SNE embedding, we flattened them to produce a (*d* = *h w c*) dimensional representation. For visualisation purposes, we colour-coded the sequences according to six expression quantiles, Q1 to Q6. In Figure S9, we show how the convolutional layers progressively group sequences of similar expressions together, with one of the strongest separations displayed by the sixth layer (Figure 7a). Overall, residual errors were evenly distributed across the sequence space, although very lowly and very highly expressed sequences displayed slightly higher error rates (Figure 7b). These results demonstrate that the model is able to progressively extract patterns in the promoter sequences that determine gene expression.

**Figure 7:**
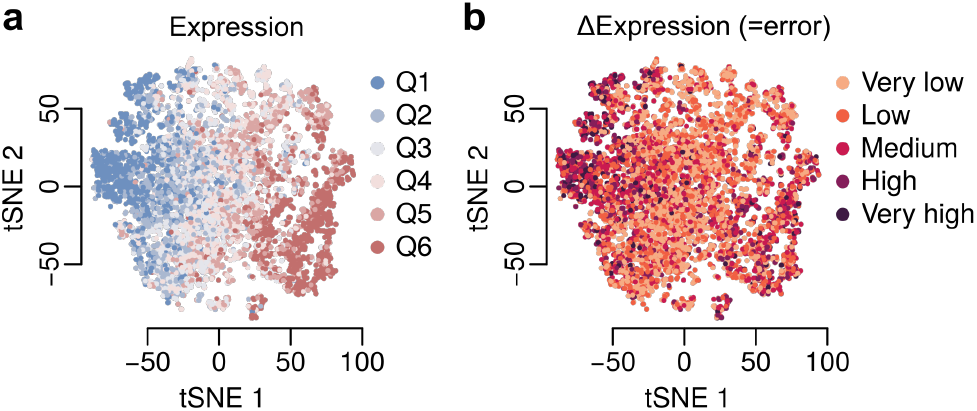
t-SNE embedding for 10,000 randomly sampled promoters from the challenge test set. The scatter plots show the two-dimensional t-SNE embeddings of the activations constructed by the last (6th) convolution layer of the Camformer model. (a) The points are coloured based on the quantile of their expression values where Q1 is the lowest and Q6 is the highest; (b) The points are coloured based on the error quantiles from very low to very high error. Embeddings for all layers are shown in Figure S9.

### 4.3 Camformer captures regulatory signals

We next asked whether it is possible to learn associations between characteristics of a promoter, such as TF binding site motifs, and the resulting expression level. Here, we utilised two in silico approaches to analyse the effect of promoter sequence on gene expression: (a) Gradient-weighted class activation map (Grad-CAM) and (b) Input perturbation analysis through in silico mutagenesis (ISM).

#### 4.3.1 Grad-CAM highlights potential regulatory hierarchy

We first used Grad-CAM to record and visualise what regions of the input promoter sequences the model attends to while making predictions. In short, Grad-CAM estimates how a perturbation in the input sequence influence the predicted output. For this experiment, we applied Grad-CAM to all 71,103 sequences in the test set. Figure S10 shows an example of a single promoter, where a strong motif is highlighted towards the 3’ end across all convolutional layers, and two additional upstream motifs are captured primarily in the deeper layers, potentially suggesting a hierarchy.

Extending this analysis to all promoters, the average saliency map per layer is shown in Figure S11. As expected, the model focuses on the variable positions 18 to 97, and pays less attention to the fixed sequences at the 5’ and 3’ ends. Moreover, the average maps indicate that the model puts higher focus towards the 3’ end of the sequence. This is consistent with a higher impact of TFs that bind near the core promoter and the transcription start site (TSS). Interestingly, the model was also strongly affected by GC content across the whole variable sequence (Figure S11). In agreement with this, we observed a strong correlation between GC content and expression (*r* = 0.263, Figure S12).

#### 4.3.2 ISM highlights transcription factor motifs

While Grad-CAM and its variants remain among the most widely used tools for interpreting deep neural networks in genomics [Zhou et al., 2023], we further conducted an in silico saturation mutagenesis experiment using input perturbation analysis [Ivanovs et al., 2021]. Here, we used sequences from the test set and systematically mutated each nucleotide across positions 18-97, skipping the flanking regions (1-17 and 98-110). This resulted in the creation of 240 new sequences—each with a different SNV—for each original sequence. We passed these to our model and recorded the predicted expression. Next, we quantified how each nucleotide in the original sequence contributes to gene expression as the negative mean per-nucleotide effect of each SNV. To find sequence motifs controlling gene expression, we calculated the entropy of the ISM sequence logo for each input promoter. Low entropy indicates high variation between nucleotide importance scores, and we hypothesised that such sequences would contain distinct regulatory motifs. Indeed, we observed that top-scoring sequences contained both activating and repressive elements, associated with high and low expression, respectively. Overall, there was a high concordance between ISM and Grad-CAM motifs, although motifs produced by ISM appears to be more well-defined (Figure S13).

### 4.4 Model predictions reveal *cis*-regulatory logic

To demonstrate how Camformer can be used to infer *cis*-regulatory logic, we next performed a SEA enrichment analysis on the training data to identify motifs that are overrepresented in high- and low-expressed sequences, respectively (Figure 8a). To reveal potential interactions, we calculated the mean expression of test set sequences for each combination of these motifs (Figure 8b). While most individual motifs displayed only small differences in expression (Figure 8b, diagonal), sequences containing the REB1 motif displayed highly elevated expression, consistent with its known role in displacing nucleosomes [Hartley and Madhani, 2009]. In contrast to individual motifs, many motif combinations resulted in substantially altered expression. To quantify this, we calculated the expression change between sequences with two motifs present together, compared to sequences containing only the first motif (Figure 8c). This revealed that most motifs have a consistently activating (REB1, ABF1, PDR3, TYE7, CBF1, TEA1, IME1, NSI1, and RAP1) or repressive (YAP7, YPR015C, NRG1, TOD6, and RPH1) effect, when present together with any other motif. In contrast, UME6 had a strongly repressive effect, only in the presence of IME1 or PDR3. This may reflect competition between TFs, or an interaction between UME6 and IME1 [Harris and Ünal, 2023].

**Figure 8:**
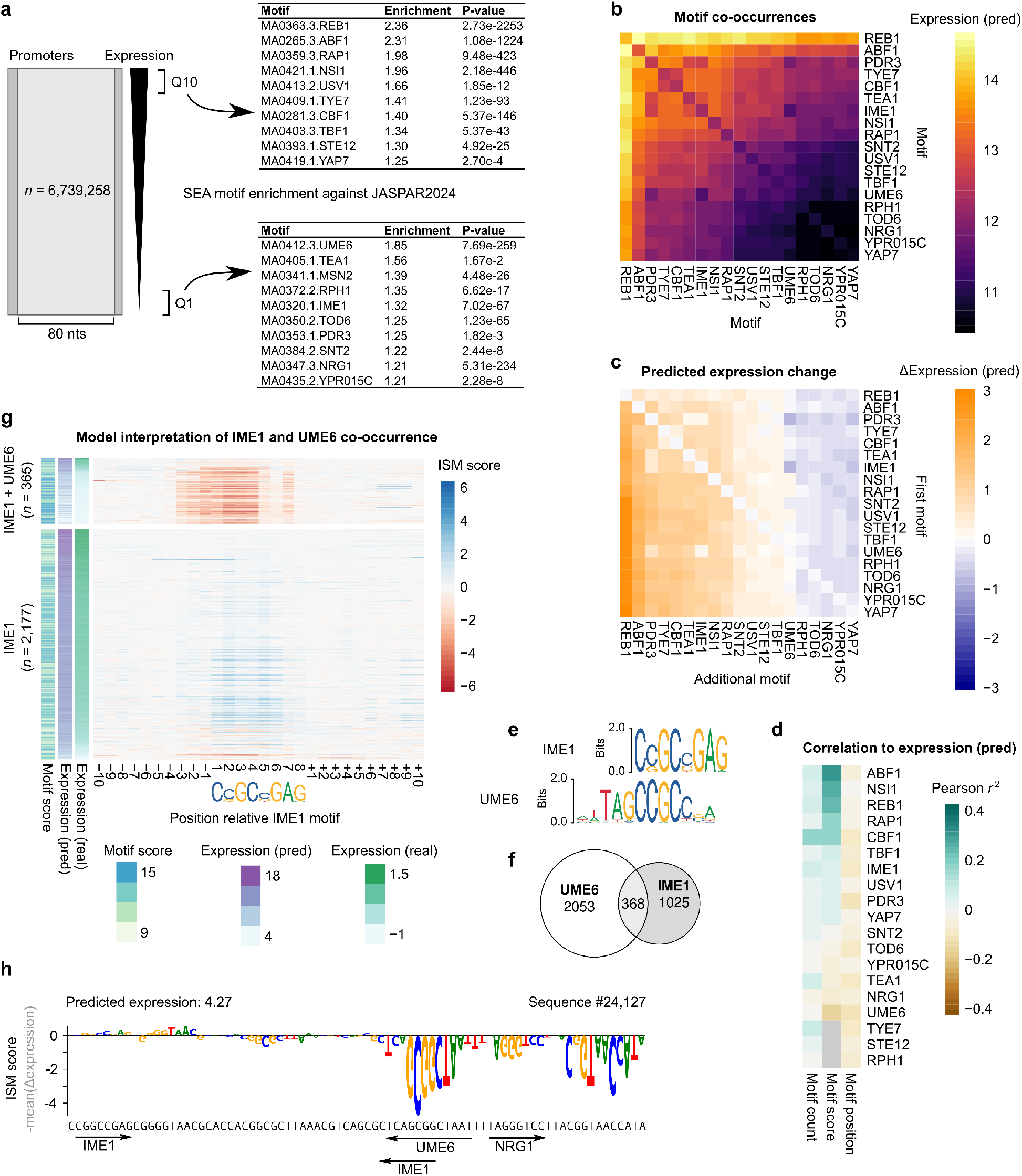
Camformer predictions reveal gene-regulatory logic. (a) The training set were divided into ten expression quantiles (Q1 to Q10) and SEA was used to identify the most enriched motifs among the highest (Q10) and lowest (Q1) expressed promoters. (b) Heatmap showing mean predicted expression for training set sequences containing pairwise combinations of selected motifs as detected by FIMO. Baseline expressions for each individual motif are shown along the diagonal. (c) Heatmap showing mean expression difference between sequences with one motif (indicated by row) and sequences with two co-occurring motifs (row and column). (d) Heatmap showing correlation between predicted expression and motif count, score, or position. Grey values indicate missing values due to all motifs having the same score. (e) JASPAR2024 motif logos show that IME1 and UME6 motifs may overlap. (f) Venn diagram illustrating that some test set sequences had both motifs. (g) Heatmap showing ISM scores per position across IME1 motifs in the test set. The IME1 motifs are grouped into those that co-occur with UME6 (*n* = 365) and those that appear on their own (*n* = 2177). (h) Example of nucleotide motifs highlighted using ISM. Positive values indicate contribution to higher expression and negative values indicate a contribution to lower expression. Positions and orientation of IME1 and UME6 motifs are shown as arrows. Only variable positions 18-97 are shown.

Each detected motif is assigned a score reflecting how well it matches its target motif. This score will roughly reflect its binding affinity and there should subsequently be a positive correlation between score and expression for activating motifs, and a negative correlation for repressive motifs. Indeed, when investigating the correlation between predicted expression and motif count, score or position (Figure 8d), we observed a high correlation for the activator ABF1 and a low correlation for UME6. Additionally, the number of motifs was generally weakly positively correlated with expression, and motif position negatively correlated, suggesting that having motifs positioned too close to the TSS may inhibit efficient transcription.

We noticed that the interaction between UME6 and IME1 could partially be explained by motif similarities (Figure 8e), with 386 sequences containing both motifs (Figure 8f). To investigate whether our model could separate the effect of IME1 on its own, or together with UME6, we grouped the IME1 motifs by whether they overlapped a UME6 motif or not (Figure 8g). Strikingly, the model consistently associated the presence of both motifs with a strong reduction in gene expression, as measured by the ISM score over the motif. In contrast, IME1 on its own had a neutral to weakly positive (Figure 8g,h). These observations suggest that models such as Camformer naturally capture and can be used to study *cis*-regulatory logic.

## 5 Discussion

In this study, we present an in-depth analysis of Camformer, a CNN-based deep learning model specifically developed to predict gene expression from promoter sequences. Despite recent advances in the application of deep learning in genomics, Camformer demonstrates that it is possible to achieve state-of-the-art performance by designing a model entirely from scratch, rather than building upon pre-existing blocks or architectures. One key advantage of this approach is the possibility to design small and efficient bespoke models such as Camformer-Mini (1.4M parameters), while maintaining competitive performance. In contrast, other top-performing submissions explored a range of recently developed architectures. For instance, LegNet used EfficientNetV2 [Tan and Le, 2021] blocks and CRMnet used a transformer-encoded U-Net structure. Notably, Proformer adopted a novel transformer architecture, featuring a Macaron-like encoder, which together with a multiple expression head strategy allowed their heavily overparameterised model (47.4M parameters) to effectively converge.

By thoroughly exploring the Camformer model design space, we show that the choice of sequence encoding and optimiser significantly impacts its performance. We systematically tested 270 model variants, but none outperformed the original model, suggesting that Camformer is located at a performance maximum. However, using an ensemble approach, we demonstrate further improvement, as previously observed with LegNet [Penzar et al., 2023]. This suggests that some residual errors could potentially be removed with a more refined model design.

Furthermore, our work demonstrates that Camformer can be used to uncover *cis*-regulatory mechanisms underlying gene regulation. For instance, using Grad-CAM and ISM, we show that Camformer recognises the effect of key TF motifs, including REB1, AZF1, UME6, and YAP7. Moreover, our model captures an interaction or hierarchy between UME6 and IME1 motifs, and can thus be used to study more complex *cis*-regulatory logic.

One particular difficulty in DREAM2022 was that while the training data consisted of random sequences, the test data sequences were not random, nor did they represent a comparable distribution of expression values. As a result, predicting gene expression on the test data was inherently difficult owing to a distribution shift [Wiles et al., 2022] between the training and test datasets. Consistent with this, all submitted models displayed strong performance on random sequences, as opposed to native sequences or sequences designed to drive unusually high or low expression levels [Rafi et al., 2024].

## Data and code availability

The DREAM challenge website is accessible at https://www.synapse.org/#!Synapse:syn28469146/wiki/617075. The associated data (training and testing) is available at https://zenodo.org/records/7395397. The code repository for this study is available at https://github.com/Bornelov-lab/Camformer.

## Acknowledgements

We are grateful to Fredrik Svensson and Maria-Anna Trapotsi who contributed to the original model as part of the Camformers team in the DREAM2022 challenge.

