## Supplementary Materials for "Predicting gene expression using millions of yeast promoters reveals *cis*-regulatory logic"

A PREPRINT

**Tirtharaj Dash**, **Susanne Bornelöv***
Cancer Research UK Cambridge Institute
University of Cambridge
Cambridge CB2 0RE
United Kingdom
{td522,smb208}@cam.ac.uk

January 25, 2025

### 1 Supplementary Methods

#### 1.1 Submission model

##### 1.1.1 Candidate model structures

We explored CNN architectures with a variable number of convolutional layers followed by flattening and up to two fully connected layers. Given that most TF motifs are only a few nts long, we reasoned that using a small kernel size might avoid overfitting. Consequently, we tested kernel sizes ranging from 3 to 15, starting with two convolutional layers and then gradually exploring deeper models, up to approximately eight layers. Since ensemble methods were not permitted, we hypothesised that a high dropout rate would achieve a similar regularisation effect.

In addition, we explored data augmentation techniques [Cao and Zhang, 2018], including the addition of reverse complement sequences, extend-and-crop techniques, and upsampling the distribution tails, but none of these augmentation techniques lead to statistically significant improvements. Similarly, we observed no improvements when introducing random noise, transforming the target value distribution, or altering the length of the promoter sequences. In contrast, introducing residual connections [Krizhevsky et al., 2012] between every other layer provided a significant improvement, and this was therefore included in the final model.

#### 1.2 Hyperparameter grid search

This section provides a detailed overview of the hyperparameter grid search. For all experiments, the training data was stratified into training (30%), validation (45%) and test (45%) (Fig. S1). The best setup of each structure was then trained on all training data using ten different random seeds.

##### 1.2.1 Structure

We considered three different CNN structures:

1. *Camformer*: The original submission model structure (as described in the main manuscript in Fig. 2b) with 16M parameters, consisting of six convolutional blocks with residual connections in alternative blocks, followed by two fully connected blocks and the output.

*corresponding author

2. *Camformer-Small*: To reduce the model's complexity (in number of parameters) drastically, we took the original model structure (above) and defined a new structure by halving the kernel sizes in the convolution blocks and the size of the two linear blocks. This approach leads to a much smaller model with approximately 3.5M parameters.
3. *Camformer-Mini*: During model design, we found a tiny model, yet it demonstrated a strong predictive performance. This structure resemble the original structure except that it does not use residual connections, different dropout rates and kernel sizes. Overall, the size of this structure is approximately 1.4M.

###### 1.2.2 Sequence encoding

A commonly used strategy for modelling genomics using CNNs is to one-hot encode each nucleotide and create a tensor representing each DNA sequence. Here, we consider some potential alternatives, which are derived using the one-hot encoding with additional information:

1. *One-hot encoding*: The classical encoding frequently used for encoding a DNA sequence. Assuming we have an alphabet $\Sigma = \{A, C, G, T\}$, the following are the encodings: $A = [1, 0, 0, 0]$, $C = [0, 1, 0, 0]$, $G = [0, 0, 1, 0]$, $T = [0, 0, 0, 1]$, and the special symbol, $N = [0, 0, 0, 0]$ denoting an unknown nucleotide.
2. *One-hot encoding with $N$ encoding*: It is easy to see that one-hot encoding treats each position (out of 4) to represent a probability, resulting in a 4-length probability vector. So, when a position in a sequence is uncertain, represented with an $N$, one can encode this as a uniform probability vector $N = [\frac{1}{4}, \frac{1}{4}, \frac{1}{4}, \frac{1}{4}]$. This encoding potentially allows some uncertainty information to be encoded.
3. *One-hot encoding with $P$ encoding*: This extends one-hot with $N$ encoding by a 5th bit to encode for whether a nucleotide is a purine (5th bit is 1) or pyrimidine (5th bit is 0). For unknown nucleotide ($N$), the 5th bit is encoded as $\frac{1}{2}$.
4. *One-hot encoding with $integer$ encoding*: Due to low sequencing coverage, 57.5% of the training data had integer expression values, indicating that the expression estimates were derived from only a small number of cells. To allow the model to use this uncertainty during training, we added a binary channel to denote whether the nucleotide originated from a sequence with integer-valued (0) or real-valued (1) expression. During inference, this bit was always set to 1. The integer encoding was inspired by LegNet [Penzar et al., 2023].
5. *One-hot encoding with both*: This encoding extends one-hot with $N$ encoding by encoding for $P$-encoding and integer encoding in its 5th and 6th channels.

###### 1.2.3 Training loss function

We study the effect of three different loss functions on the model's performance:

1. *$L_1$ loss*: It is computed as the absolute difference between true expression value $y$ and predicted expression value $\hat{y}$. That is, $L_1(y, \hat{y}) = |y - \hat{y}|$. $L_1$ is less sensitive to outliers and noisy data.
2. *$L_2$ loss*: It is also called the squared error and is computed as $L_2(y, \hat{y}) = (y - \hat{y})^2$. $L_2$ loss penalises outliers and noise in the data.
3. *Huber loss*: Huber loss [Huber, 1992] utilises the benefits of both $L_1$ loss and $L_2$ loss. It results in a squared term if the absolute element-wise error falls below some threshold $\delta$ and a $\delta$-scaled $L_1$ term otherwise, as shown in the scalar-form below. We used $\delta = 0.9$ in all our experiments.

$$L_H(y, \hat{y}) = \begin{cases} \frac{1}{2}(y - \hat{y})^2, & \text{if } |y - \hat{y}| < \delta \\ \delta\left(|y - \hat{y}| - \frac{\delta}{2}\right), & \text{otherwise} \end{cases}$$

We use the mean of these losses over a batch of promoter sequences during training.

###### 1.2.4 Optimiser

We use two variants of the stochastic gradient descent (SGD) optimisation algorithm:

1. *AdamW*: Adam optimiser [Kingma and Ba, 2015] with weight decay [Loshchilov and Hutter, 2019], which improves parameter regularisation and prevents over-fitting during training.
2. *Lion*: Evolved signed momentum [Chen et al., 2024], a potentially more stable version over standard SGD algorithms, and suited for larger batch sizes due to the nature of the parameter update. It combines the principles of signed gradient updates with momentum, which significantly reduces memory usage and enhances computational efficiency.

###### 1.2.5 LR scheduler

For SGD-based optimisers, LR plays a crucial role in model training, contributing significantly to the stability of the training process. Learning rate scheduling is often considered a good practice for an automated adaptation of a suitable LR during the training itself. Here, we study three different strategies for LR scheduling: (1) No explicit scheduler, (2) One-cycle LR [Smith and Topin, 2019] as used by the 1st and 3rd ranked submissions [Rafi et al., 2024], and (3) Reduce LR on training plateau.

**Model search space**

The settings described above were used to systematically construct variants of the Camformer model using grid-search: That is, training and evaluating each model individually on a validation set. Overall, the grid search space consists of 270 models (3 different CNN structures and $5 \times 3 \times 2 \times 3 = 90$ different sets of hyperparameters).

**Training hyperparameters**

For training the Camformer models, we use randomly drawn 90% of the training dataset (≈5.8M). We use a batch size of 256 and learning rate of 0.001 for AdamW optimiser. The patience period is set to 10 for early-stopping. We use the sum of Pearson and Spearman correlation coefficients, $r + \rho$ as validation objective function during our training. The Large variant of the Camformer model takes around 6 hours in the Dell workstation with 64GB main memory and NVIDIA GeForce RTX 4090 (with 24GB GPU memory). The final Camformer codebase was developed using PyTorch.

##### 1.3 Interpretability Tools

We implement Grad-CAM [Selvaraju et al., 2017] and in silico mutagenesis (ISM) in our explanatory analysis of the Camformer models.

###### 1.3.1 Grad-CAM

Grad-CAM highlights important regions in the input sequence by analysing the gradients of the model's output with respect to the inputs received by each convolutional layer. It highlights which parts of the input sequences contribute most to the decision process of the CNN. This method evaluates the sensitivity of the model to different regions of the promoter sequence. In our experiments, a one-hot encoded promoter sequence ($\mathbf{X}$) is passed as input to Camformer. The gradient of the output is computed with respect to the input, $\mathbf{X}$, as $\frac{\partial \hat{y}}{\partial \mathbf{X}}$, and a feature map for each layer is collected. Finally, Grad-CAM combines the mean of the feature maps with the gradients to determine how important each part of the input is for the model's decision. In essence, the gradients at the input layer indicate how sensitive the model's output is to changes in the input ($0 \rightarrow 1$ or $1 \rightarrow 0$). Internally, the gradients are distributed across all positions, reflecting the network's weights and activations, which collectively determine the influence of each input feature on the output expression.

###### 1.3.2 In silico mutagenesis (ISM)

ISM is a simple approach to observe how the model processes and responds to variations in the input sequence. While Grad-CAM allows us to probe the model in an automated manner by using the output gradients at every layer, ISM allows a more systematic study of position-wise effect on the model's output without having to look at every layer's internal representations. It involves systematically mutating each position of the input sequence and observing the change in the model's output. This method evaluates the sensitivity of the model to each nucleotide at each position. In our implementation, we compute an average effect of mutations (A → {C, G, T}, C → {A, T, G}, . . . ) by observing the change in the predicted expression with respect to the prediction without the mutation. We then visualise these quantitative mutation-induced effects on a sequence logo plot.

To quantify goodness of ISM and to help us highlight the distinctive motifs in a sequences, we compute an entropy measure over the change in expression values upon single-nucleotide in-silico mutations. A high entropy value would then mean a sequence where all the positions are equally important and ISM couldn't identify a continuous sub-sequence that is influential in prediction. In contrast, the opposite is true for the lowest entropy value, that is, a sequence does contain one or more clearly identifiable motifs. The whole process is illustrated as below.

Let $S$ be an input sequence with predicted expression $e$. Let $Mut(S)$ be the set of all possible mutations on $S$ at all positions in the range $[17, 96]$ (80 nts) and $\Delta(e)$ be the set of changes in expression values due to the individual mutations. We first calculate a position-specific scoring matrix (PSSM) to record mutation-induced expression change due to each position, denoted by $PSSM(S, i, x)$, defined as the mean $\Delta(e)$ at position $i$ due to change of the original nucleotide to $x$ in sequence $S$. Then, the entropy of the sequence $S$, denoted by $H(S)$ is:

$$\texttt{Importance}(S,i) = \sum_{x\in\{A,C,G,T\}} PSSM(S,i,x)$$

$$\texttt{prob}(S,i) = \frac{|\texttt{Importance}(S,i)|}{\sum_j |\texttt{Importance}(S,j)|}$$

$$H(S) = -\sum_i \texttt{prob}(S,i) \cdot \log_2(\texttt{prob}(S,i))$$

It is easy to see that $H(S)$ is the maximum if $\texttt{prob}(S,i) = \texttt{length}(S)^{-1}$, $\forall i$.

##### 1.4 Sequence Analysis using Bioinformatics Tools

We use MEME Suite [Bailey et al., 2015] (version 5.5.7) for analysing the promoter sequences obtained from the expression quantiles and ISM computation. We use the following tools along with their default parameter settings within the MEME suite: (1) Simple Enrichment Analysis (SEA [Bailey and Grant, 2021]) for motif discovery and (2) Finding individual motifs occurences (FIMO [Grant et al., 2011]) for finding positions of a motif in a sequence. For completeness, some important parameters that we used in these tools are as follows: SEA($m$-order shuffled sequences as control (negative), $m = 2$; `thresh`: enrichment $E$-values up to 10); FIMO(`thresh`: $p$=1e-4). For FIMO, we use a set of well-characterised TF motifs from *S. cerevisiae*, which is publicly available at the JASPAR2024 database [Rauluseviciute et al., 2024]. There were a total of 170 motifs used for the analysis, as listed (including their versions) in Table **S1**.

##### 1.5 Use of large language models

Parts of the manuscript was edited with assistance of ChatGPT using prompts of the type "Please provide suggestions for how to improve the writing of the following section, including a detailed explanation of each change" to improve grammar and clarity.

| | | | |
|---|---|---|---|
| MA0271.1.ARG80 | MA0387.1.SPT2 | MA0372.2.RPH1 | MA0436.2.YPR022C |
| MA0274.1.ARR1 | MA0388.1.SPT23 | MA0374.2.RSC3 | MA0438.2.YRM1 |
| MA0275.1.ASG1 | MA0391.1.STB4 | MA0375.2.RSC30 | MA0439.2.YRR1 |
| MA0276.1.ASH1 | MA0392.1.STB5 | MA0376.2.RTG3 | MA0441.2.ZMS1 |
| MA0277.1.AZF1 | MA0393.1.STE12 | MA0377.2.SFL1 | MA0266.2.ABF2 |
| MA0283.1.CHA4 | MA0394.1.STP1 | MA0378.2.SFP1 | MA0272.2.ARG81 |
| MA0287.1.CUP2 | MA0399.1.SUT1 | MA0380.2.SIP4 | MA0273.2.ARO80 |
| MA0288.1.CUP9 | MA0401.1.SWI4 | MA0381.2.SKN7 | MA0267.2.ACE2 |
| MA0289.1.DAL80 | MA0404.1.TBS1 | MA0382.3.SKO1 | MA0278.2.BAS1 |
| MA0290.1.DAL81 | MA0405.1.TEA1 | MA0384.2.SNT2 | MA0279.3.CAD1 |
| MA0291.1.DAL82 | MA0407.1.THI2 | MA0385.2.SOK2 | MA0280.2.CAT8 |
| MA0297.1.FKH2 | MA0409.1.TYE7 | MA0386.2.SPT15 | MA0281.3.CBF1 |
| MA0299.1.GAL4 | MA0417.1.YAP5 | MA0389.2.SRD1 | MA0282.2.CEP3 |
| MA0304.1.GCR1 | MA0419.1.YAP7 | MA0390.2.STB3 | MA0284.3.CIN5 |
| MA0307.1.GLN3 | MA0421.1.NSI1 | MA0395.2.STP2 | MA0285.2.CRZ1 |
| MA0311.1.HAL9 | MA0426.1.YHP1 | MA0396.2.STP3 | MA0286.2.CST6 |
| MA0313.1.HAP2 | MA0429.1.YLL054C | MA0397.2.STP4 | MA0351.2.DOT6 |
| MA0318.1.HMRA2 | MA0432.1.YNR063W | MA0398.2.SUM1 | MA0292.2.ECM22 |
| MA0320.1.IME1 | MA0437.1.YPR196W | MA0400.2.SUT2 | MA0293.2.ECM23 |
| MA0321.1.INO2 | MA0440.1.ZAP1 | MA0402.2.SWI5 | MA0294.2.EDS1 |
| MA0322.1.INO4 | MA0329.2.MBP1 | MA0403.3.TBF1 | MA0420.2.ERT1 |
| MA0323.1.IXR1 | MA0333.2.MET31 | MA0431.2.TDA9 | MA0295.2.FHL1 |
| MA0324.1.LEU3 | MA0334.2.MET32 | MA0406.2.TEC1 | MA0296.2.FKH1 |
| MA0326.1.MAC1 | MA0336.2.MGA1 | MA0350.2.TOD6 | MA0268.2.ADR1 |
| MA0327.1.HMRA1 | MA0337.2.MIG1 | MA0265.3.ABF1 | MA0300.2.GAT1 |
| MA0328.1.MATALPHA2 | MA0338.2.MIG2 | MA0408.2.TOS8 | MA0301.2.GAT3 |
| MA0330.1.MBP1::SWI6 | MA0339.2.MIG3 | MA0410.2.UGA3 | MA0302.2.GAT4 |
| MA0331.1.MCM1 | MA0343.2.NDT80 | MA0412.3.UME6 | MA0303.3.GCN4 |
| MA0332.1.MET28 | MA0347.3.NRG1 | MA0411.2.UPC2 | MA0305.2.GCR2 |
| MA0340.1.MOT3 | MA0348.2.OAF1 | MA0422.2.URC2 | MA0306.2.GIS1 |
| MA0341.1.MSN2 | MA0349.2.OPI1 | MA0413.2.USV1 | MA0308.2.GSM1 |
| MA0342.1.MSN4 | MA0352.3.PDR1 | MA0414.2.XBP1 | MA0309.2.GZF3 |
| MA0344.1.NHP10 | MA0354.2.PDR8 | MA0415.2.YAP1 | MA0269.2.AFT1 |
| MA0353.1.PDR3 | MA0355.2.PHD1 | MA0416.2.YAP3 | MA0310.2.HAC1 |
| MA0356.1.PHO2 | MA0358.2.PUT3 | MA0418.2.YAP6 | MA0312.3.HAP1 |
| MA0357.1.PHO4 | MA0359.3.RAP1 | MA0423.2.YER130C | MA0314.3.HAP3 |
| MA0364.1.REI1 | MA0360.2.RDR1 | MA0424.2.YER184C | MA0316.2.HAP5 |
| MA0366.1.RGM1 | MA0361.2.RDS1 | MA0425.2.YGR067C | MA0317.2.HCM1 |
| MA0368.1.RIM101 | MA0362.2.RDS2 | MA0428.2.YKL222C | MA0270.2.AFT2 |
| MA0370.1.RME1 | MA0363.3.REB1 | MA0430.2.YLR278C | MA0319.2.HSF1 |
| MA0371.1.ROX1 | MA0365.2.RFX1 | MA0433.2.YOX1 | MA0325.2.LYS14 |
| MA0373.1.RPN4 | MA0367.2.RGT1 | MA0434.2.YPR013C | |
| MA0379.1.MOT2 | MA0369.2.RLM1 | MA0435.2.YPR015C | |

**Table S1:** JASPAR database of 170 transcription factor motifs in *S. cerevisiae*.

#### 2 Supplementary Figures

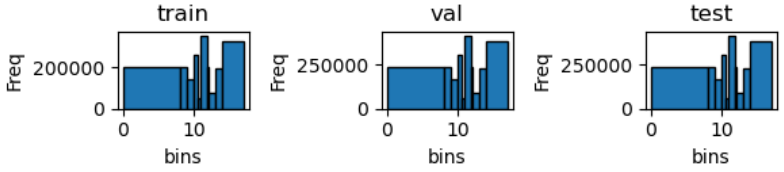


**Figure S1:** Stratified distribution of YFP expressions for model tuning. For the systematic grid search of Camformer model construction, the official DREAM challenge dataset ($n = 6.74$M) was stratified and split into training (30%) and the rest (70%) was split equally between validation and testing. The histogram of these three splits is shown below by binning the expression quantiles.

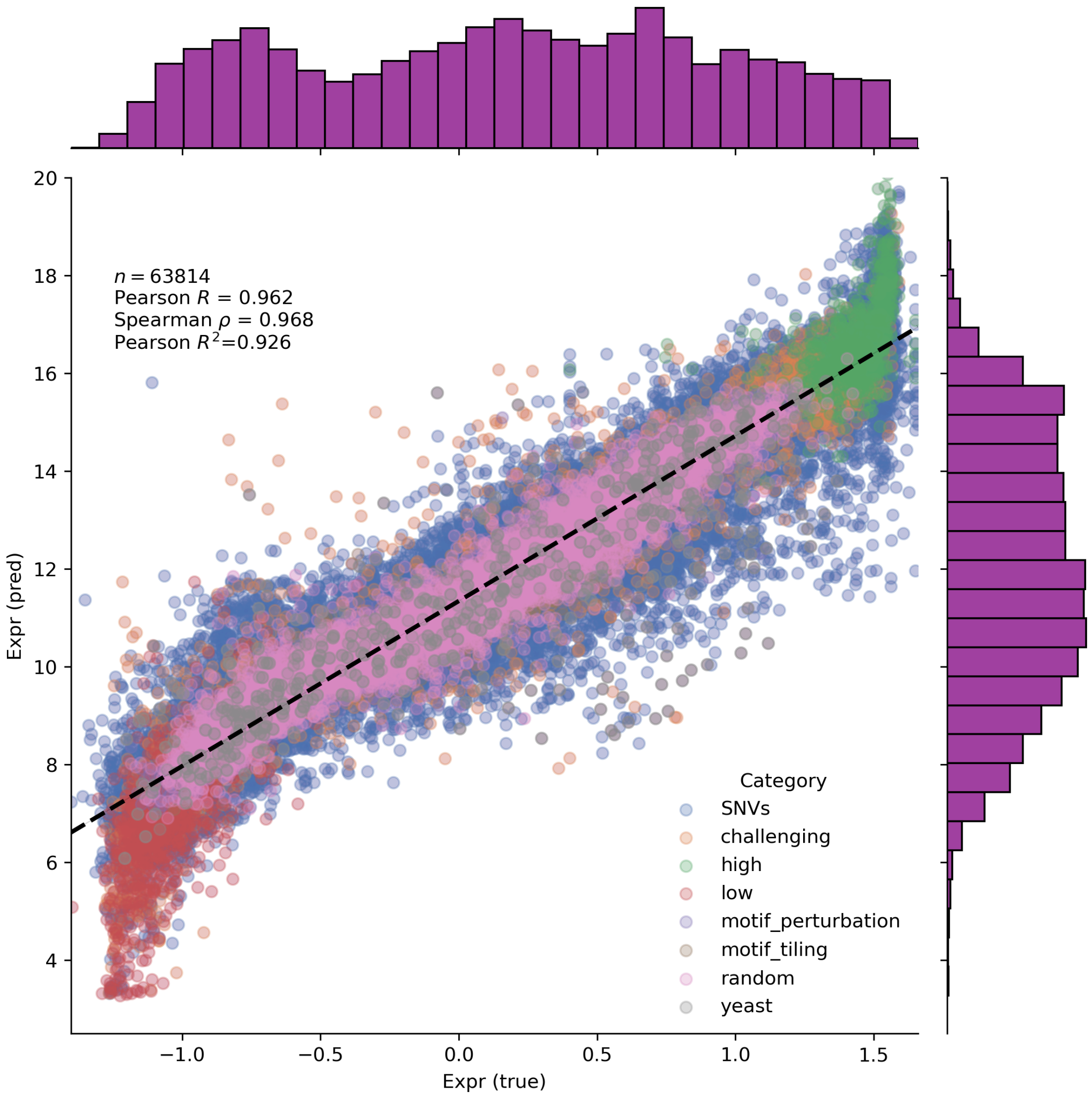


**Figure S2:** Predictive performance of the Camformer (original) model on the private split of the official DREAM challenge dataset. Each category of the promoter sequence is colour-coded. This figure helps visualise the categories of the sequences for which the Camformer model performs better or worse.

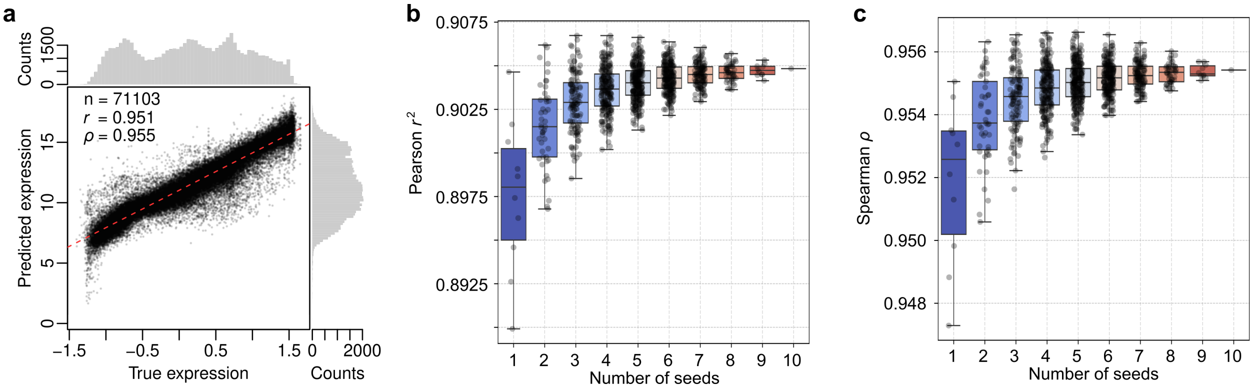


**Figure S3:** Camformer-Mini predicts gene expression from promoter sequence. The model was evaluated on the DREAM2022 test dataset ($n = 71,103$). (a) Scatter plot between true and predicted expression values, (b) Predictive performance measured using Pearson $r^2$ across ten individual replicates and ensemble models constructed by taking the mean prediction from $k$ replicates (where $k \in \{2, \ldots, 10\}$). A replicate here refers to a model constructed using a random seed, determining the initialisation of model parameters (weights) before training. (c) Same as b), but showing Spearman $\rho$. Please note that the range of expression values in the test dataset differed from that of the training dataset.

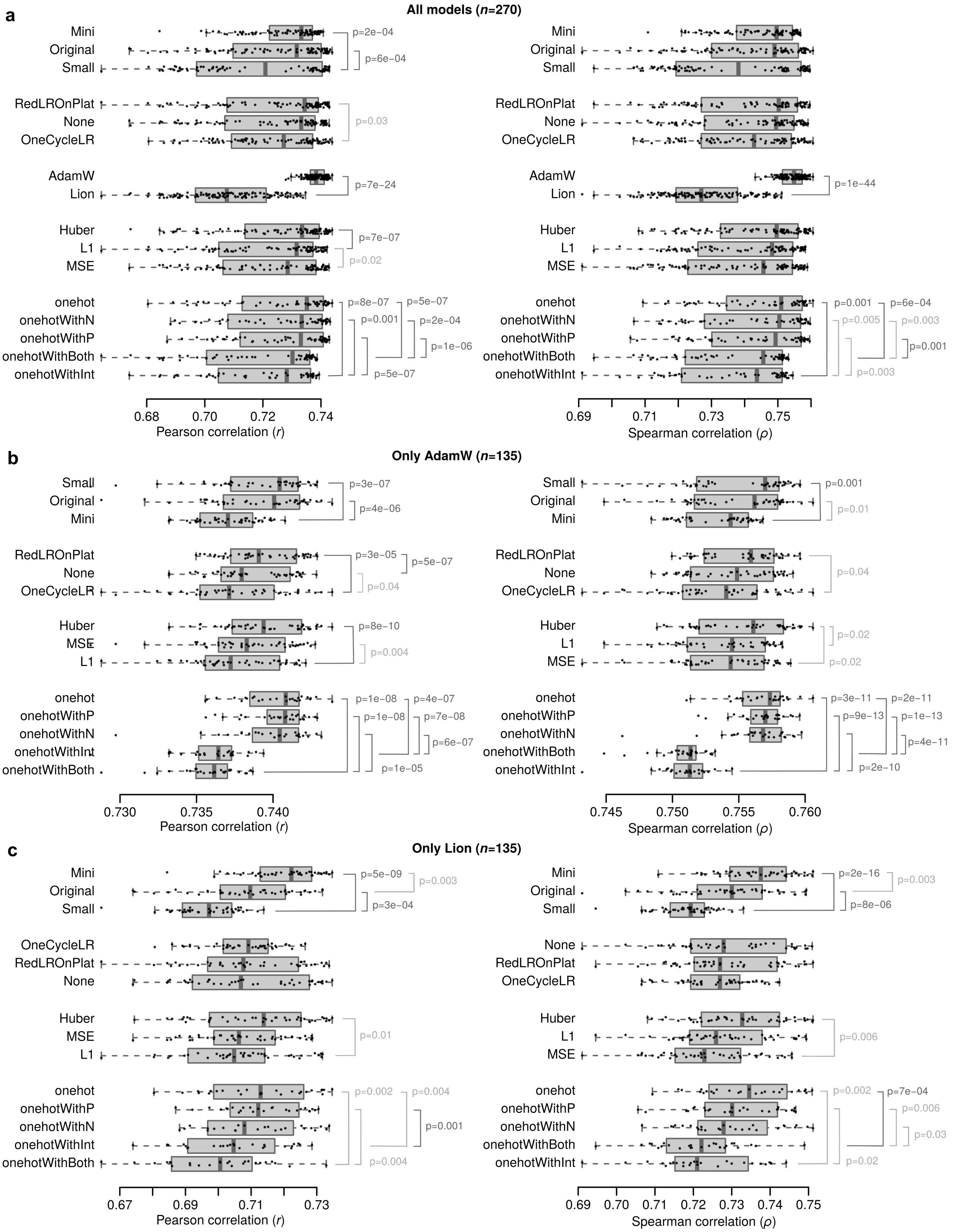


**Figure S4:** Overview of the hyperparameter grid search. (a) Performance of all 270 models across architecture (Small, Original, Mini), scheduler (ReduceLROnPlateu, None, OneCycleLR), optimiser (AdamW, Lion), loss (Huber, MSE, L1), and encoding (onehot, onehotWithP, onehotWithN, onehotwithInt, onehotWithBoth), respectively; (b) Same as a), but only for 135 models trained with AdamW; (c) Same as a), but only for 135 models trained with Lion. Statistical significance was assessed using a two-sided Wilcoxon Rank Sum test. Differences that did not pass Bonferroni correction for multiple testing (40 tests; requiring $p < 0.00125$) are indicated in lighter gray.

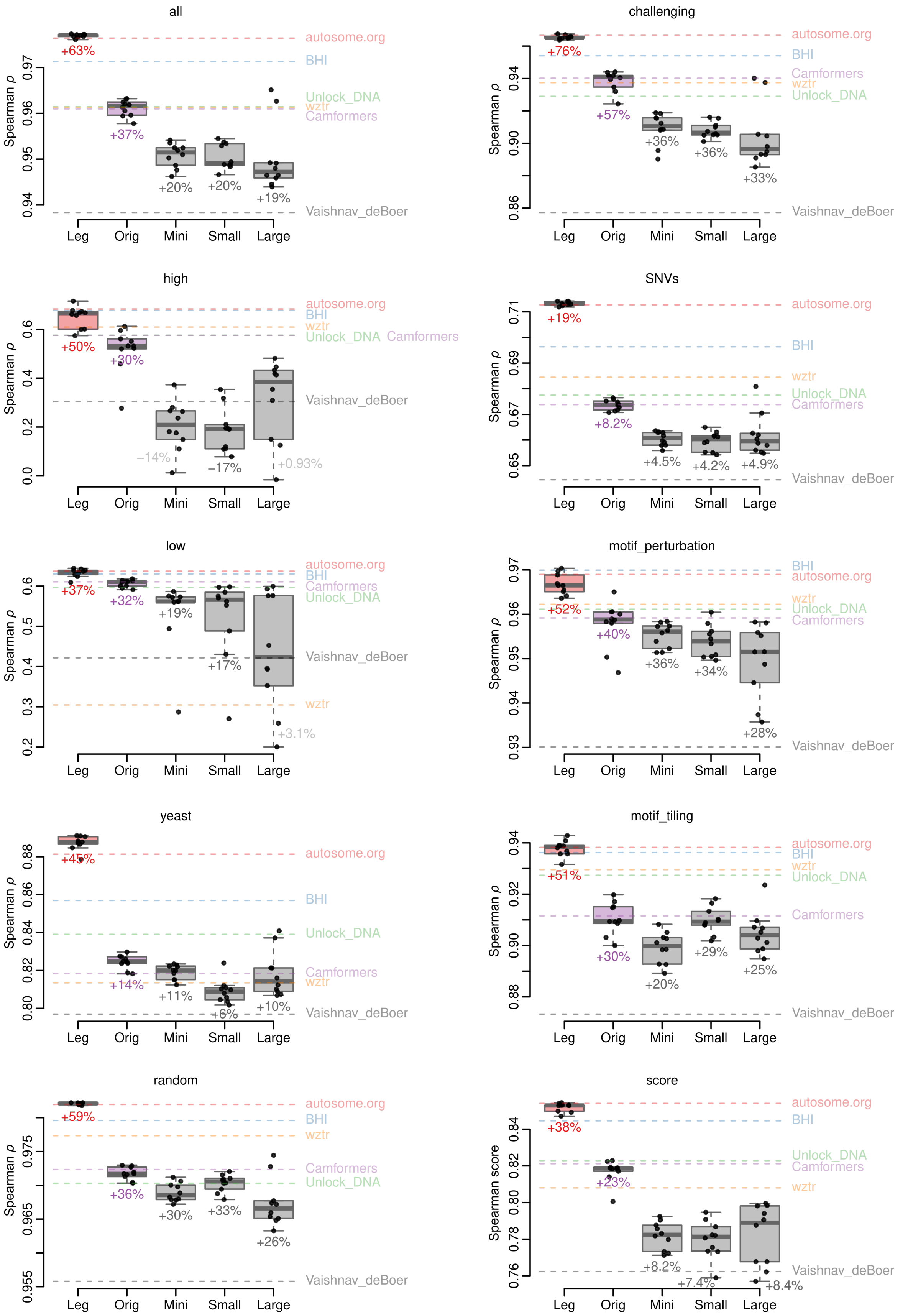


**Figure S5:** Comparison showing the predictive performance of different variants of Camformer models (Original, Large, Small and Mini) and LegNet (Leg) across 10 replicates. The box-plots show the Spearman correlation coefficients between predicted and true expression for all sequences in the private test set of the DREAM challenge, the eight different distinct subsets of evaluation sequences, and the combined evaluation score. The annotations on the figure indicate baseline-normalised improvement against Vaishnav_deBoer (gray dashed line). Improvements that were not significantly different to the baseline (two-sided Student's t-test) are shown in a more transparent colour.

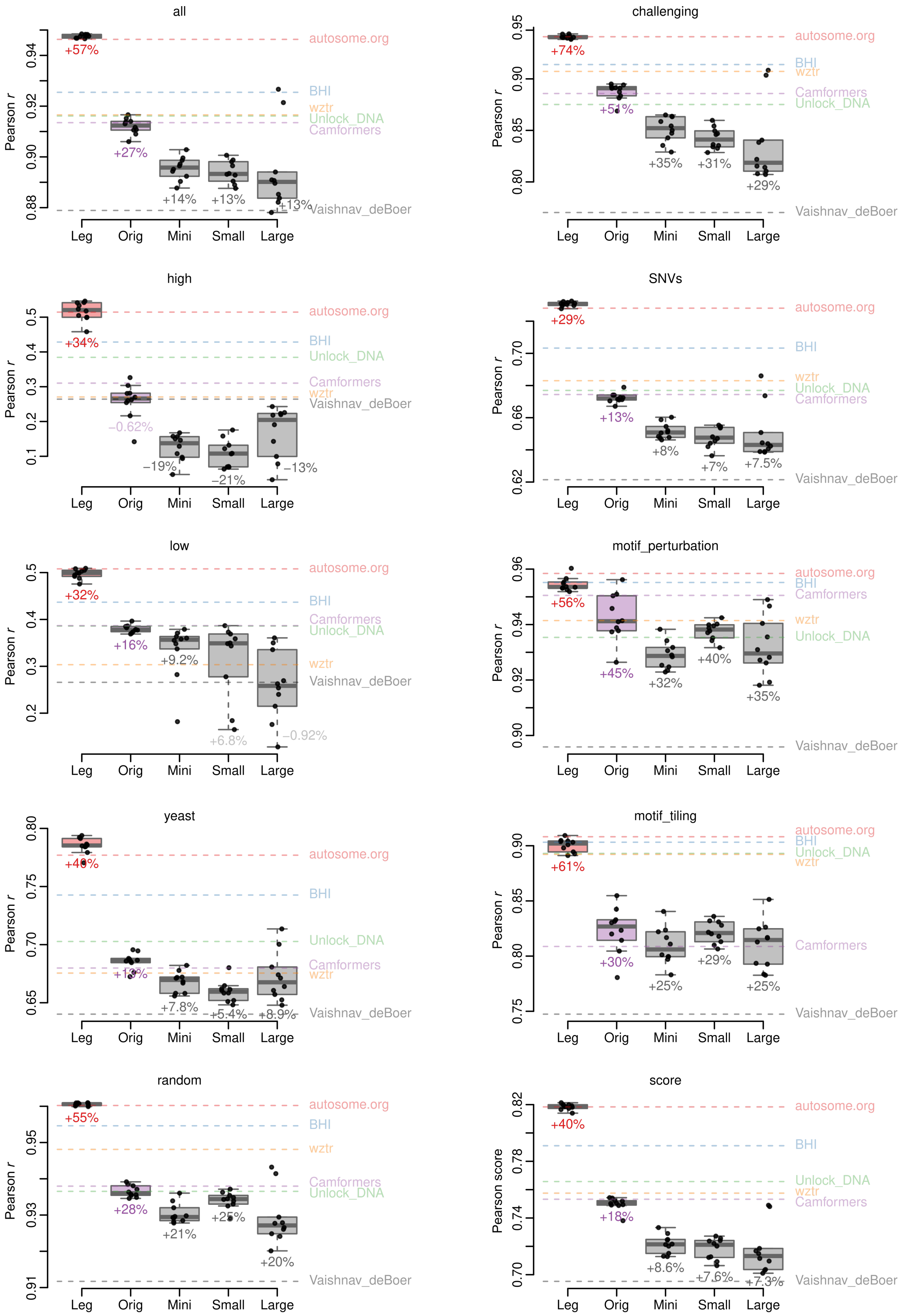


**Figure S6:** Comparison showing the predictive performance of different variants of Camformer models and LegNet (Leg) across 10 replicates. The box plots show the Pearson correlation coefficients between predicted and true expression for all sequences in the private test set of the DREAM challenge, the eight different distinct subsets of evaluation sequences, and the combined evaluation score. The annotations on the figure indicate baseline-normalised improvement against Vaishnav_deBoer (gray dashed line). Improvements that were not significantly different to the baseline (two-sided Student's t-test) are shown in a more transparent colour.

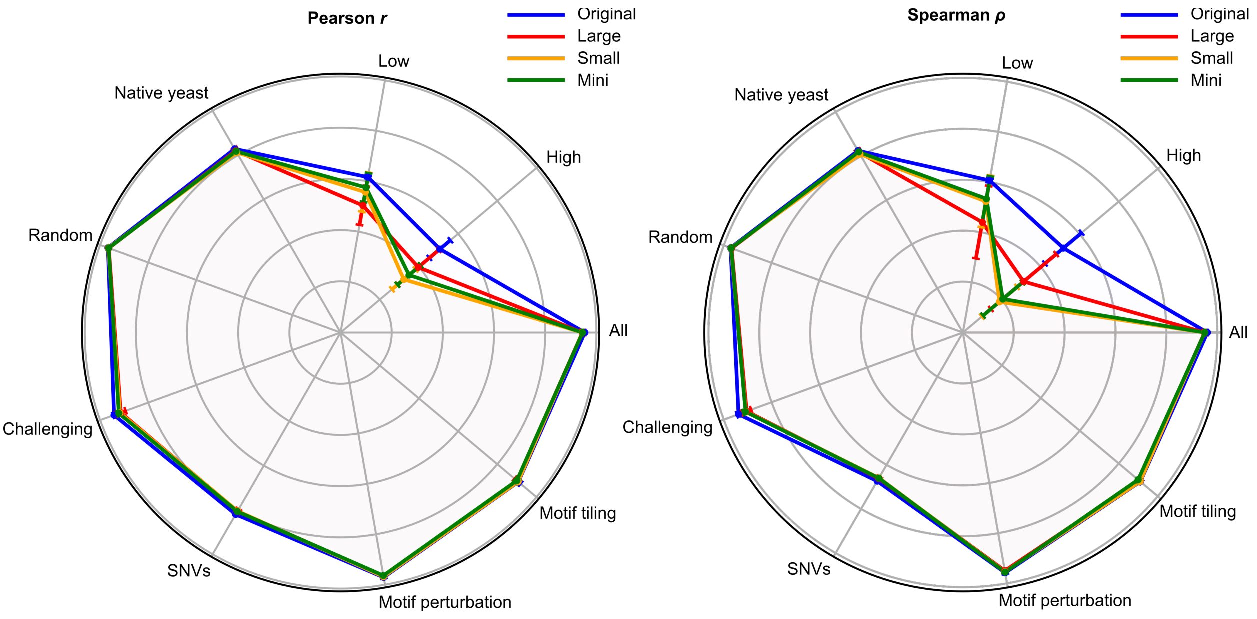


**Figure S7:** Comparison showing the predictive performance of different variants of Camformer models (Original, Large, Small and Mini) using a radar plot. This figure provides a condensed visualisation of Figure S5 and Figure S6. Left: Pearson $r$, Right: Spearman $\rho$. The figure suggests a superior performance of the original variant as compared to that of the other Camformer variants.

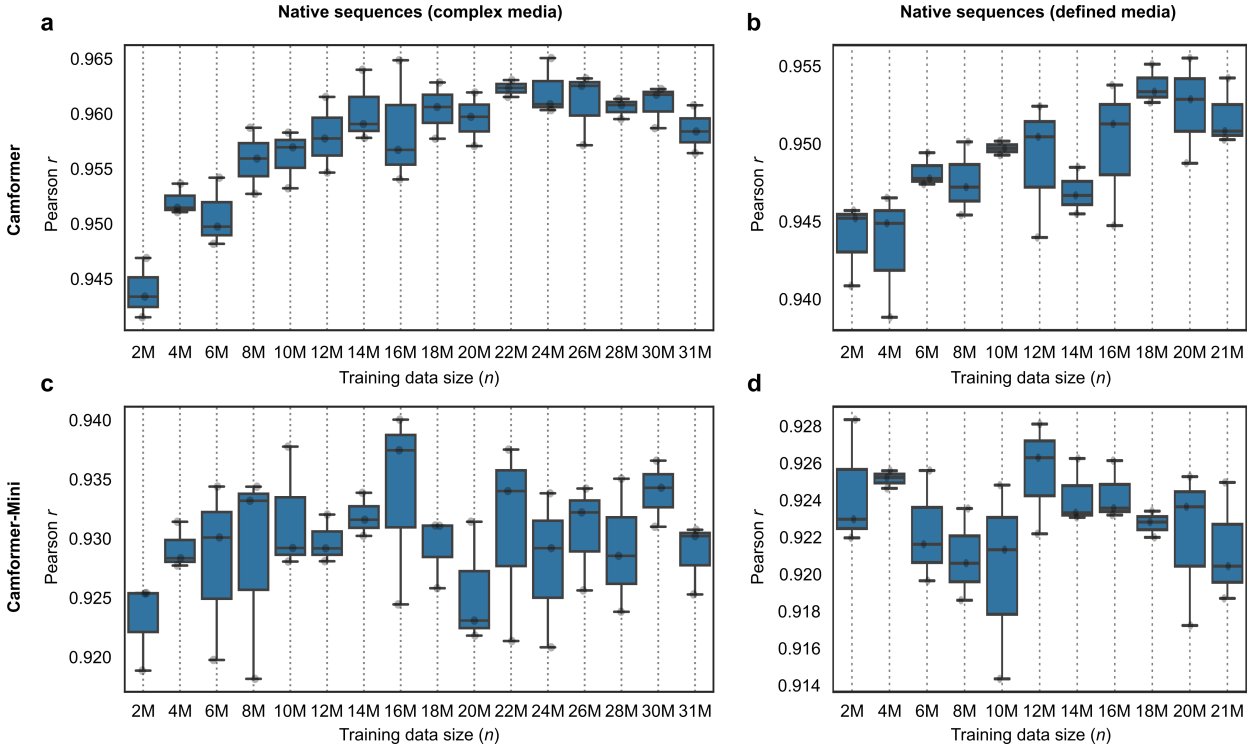


**Figure S8:** Box plots showing model performance as a function of training data size. Performance shown as Pearson $r$ for (a,b) Camformer or (c,d) Camformer-Mini models trained on native promoter activity measured in (a,c) complex (YPD) or (b,d) defined (SD-Ura) growth media. The models were trained with increasing 2M chunks (2M, 4M, ...) of the training data and their performances were measured on its corresponding test sets (complex medium: $n = 3929$; defined medium: $n = 3978$). Each model was trained three times using different random seeds.

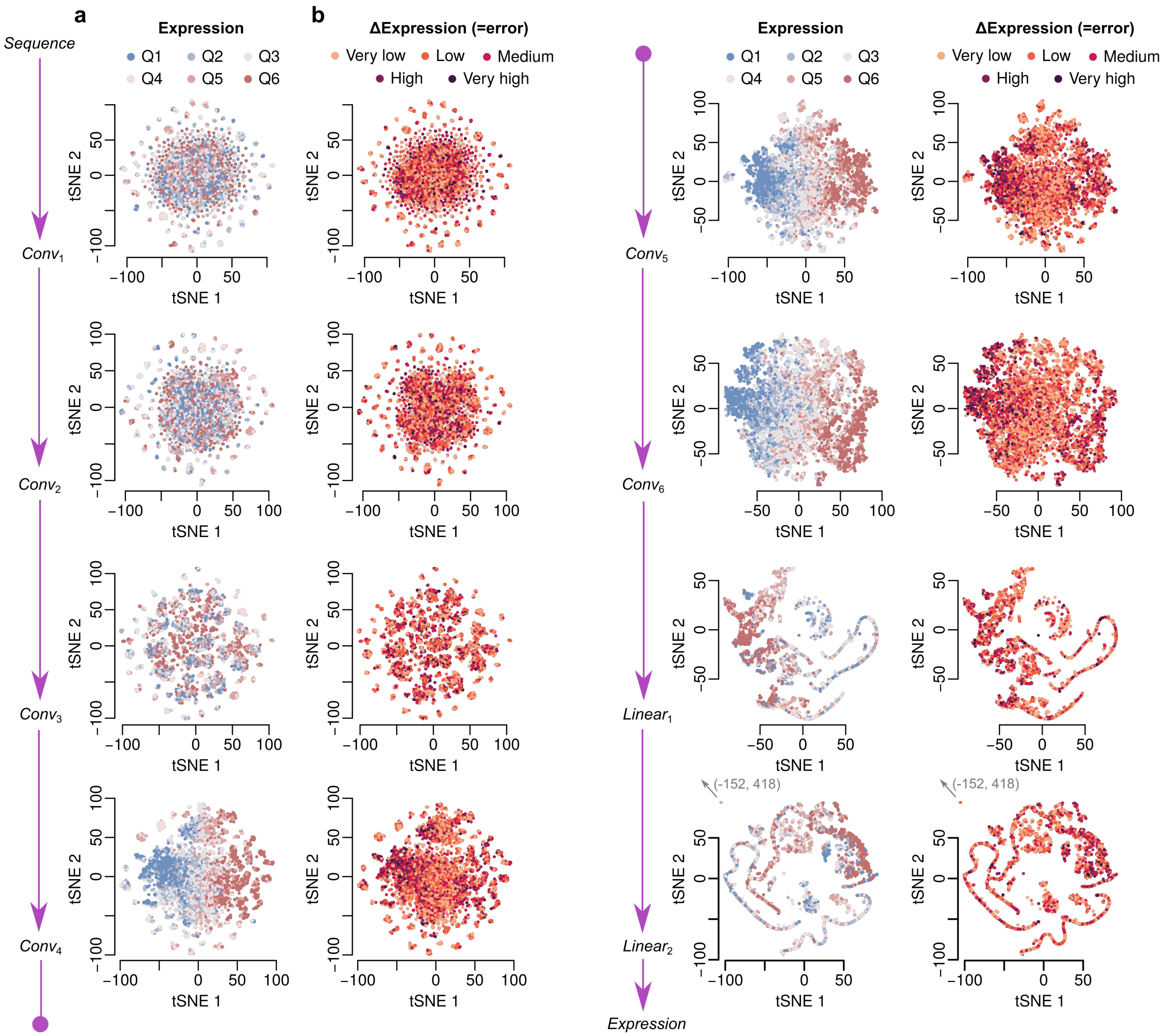


**Figure S9:** t-SNE embedding space for randomly sampled 10,000 promoter sequences from the challenge testing dataset. The scatter plots show the two-dimensional t-SNE embeddings of the activations constructed by the convolutional layers ($Conv_1$ through to $Conv_6$) followed by the linear layers ($Linear_1$ and $Linear_2$) of the Camformer model. (a) The points are coloured based on the quantile of their expression values where Q1 is the Lowest and Q6 is the Highest; (b) The points are coloured based on the error quantiles from Very low error to Very high error. The figure shows how the model progressively refines the representation of the promoter sequences and starts separating them based on the properties that define their expression levels as they move through different layers. The position of an outlier data point in $Linear_2$ is indicated with an arrow.

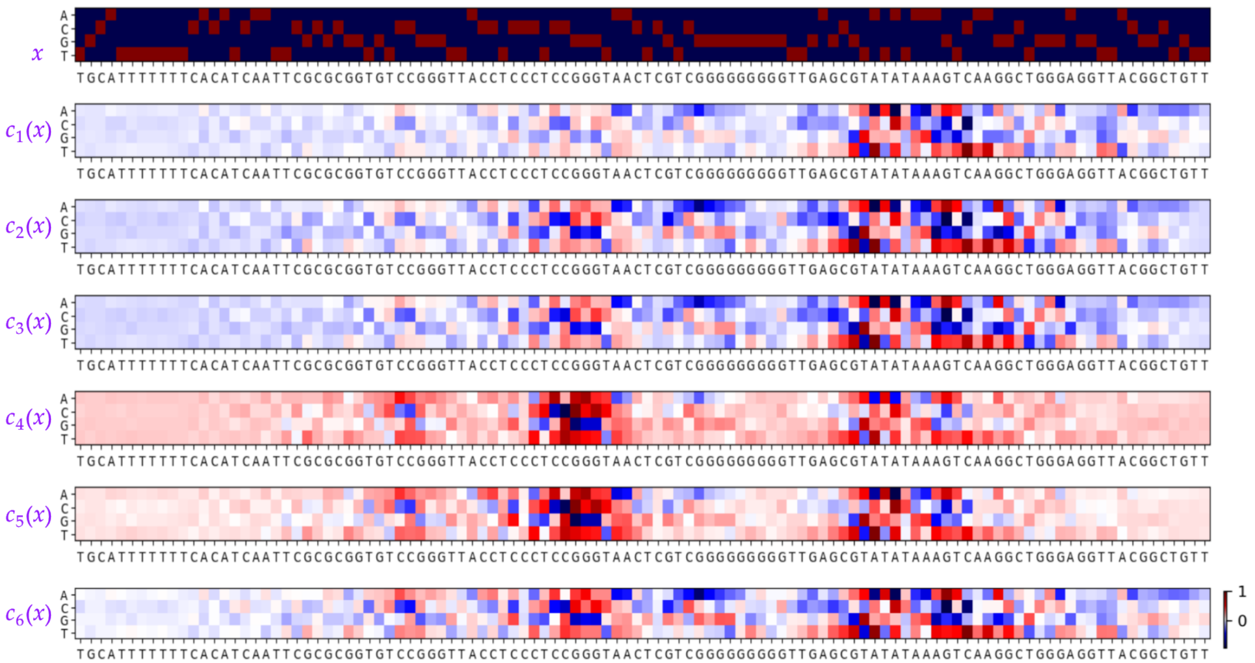


**Figure S10:** Grad-CAM saliency heatmaps for a single promoter sequence showing how the Camformer model takes the one-hot encoded sequence $x$ and the subsequent convolution layers ($c_1$ through to $c_6$) start focusing their attention on several regions (potential motifs) in the promoter, allowing the model in the end to predict an expression level. Here $c_i(x)$ is used to 'loosely' denote a transformation of the input sequence by the convolution layer $c_i$. It is clear that each layer primarily focus on the 80 nt variable region between positions 18-97 and on the flanking sequence at the 3' end, and that much less focus is on the 17 nt flanking sequences at the 5' end. A more comprehensive visualisation is shown in Figure S11.

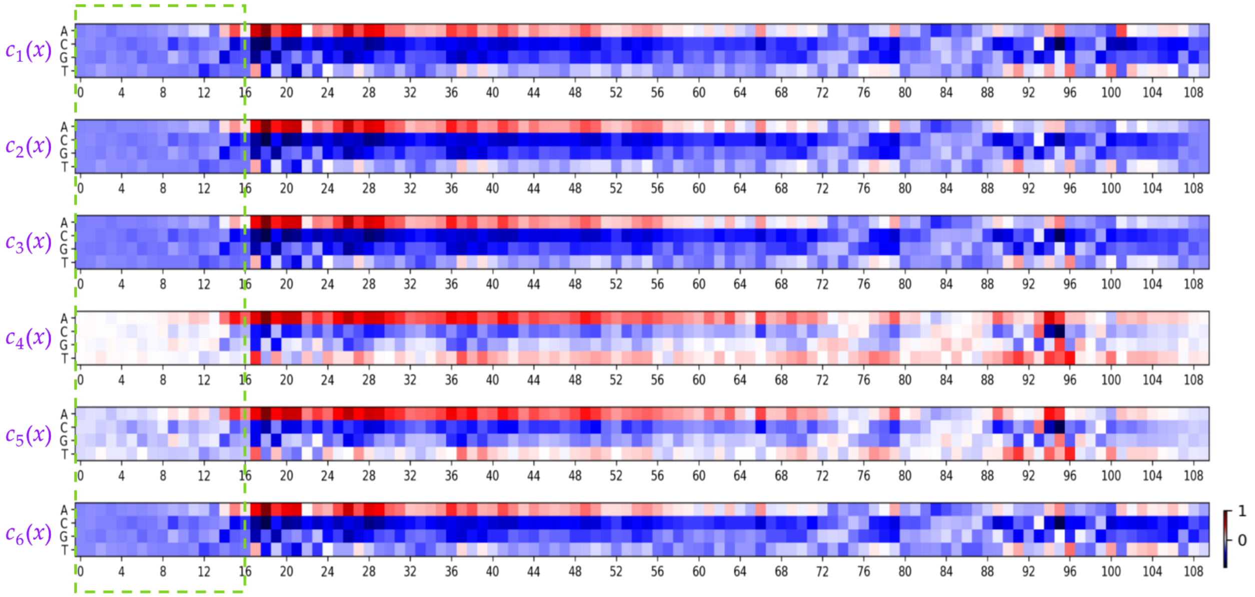


**Figure S11:** Average Grad-CAM saliency heatmaps for all the promoter sequences in the challenge test set (n=71,103). Each convolution layer ($c_i$) primarily focus only on the 80 nt variable sequence between positions 18-97 and on the flanking sequence at the 3' end, and that much less focus is on the 17 nt flanking sequences at the 5' end (green dashed area).

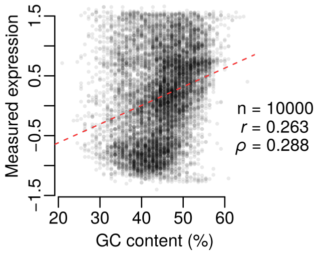


**Figure S12:** Effect of nucleotide composition on expression values of the promoter sequences. Scatter plot between GC content and gene expression. The results are based on 10,000 randomly sampled sequences from the challenge test set.

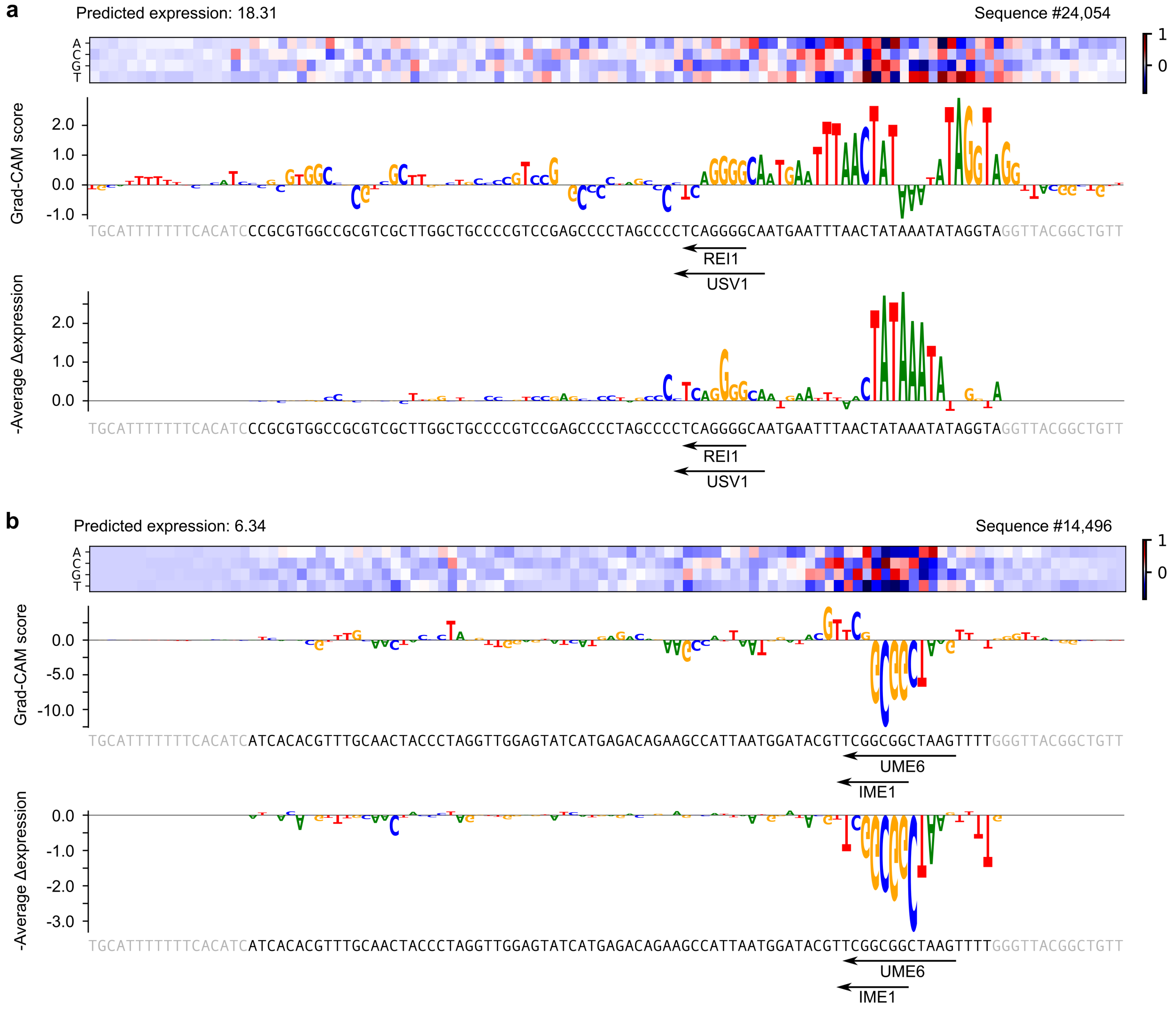


**Figure S13:** Examples of sequence motifs highlighted using Grad-CAM and ISM. Shown here is a sequence with (a) high or (b) low expression from the challenge test set. A Grad-CAM saliency map was obtained from the last convolutional layer of Camformer (top) and was visualised using LogoMaker (middle). For the same sequence, we used ISM to identify the exact nucleotide positions that are crucial for the model's prediction (bottom).

Alex Krizhevsky, Ilya Sutskever, and Geoffrey E Hinton. Imagenet classification with deep convolutional neural networks. *Adv Neural Inf Process Syst*, 25, 2012.

Ilya Loshchilov and Frank Hutter. Decoupled weight decay regularization. In *Int Conf Learn Represent*, 2019.

Dmitry Penzar, Daria Nogina, Elizaveta Noskova, et al. Legnet: a best-in-class deep learning model for short dna regulatory regions. *Bioinformatics*, 39(8):btad457, 2023.

Abdul Muntakim Rafi, Daria Nogina, Dmitry Penzar, et al. A community effort to optimize sequence-based deep learning models of gene regulation. *Nat Biotechnol*, pages 1–11, 2024.

Ieva Rauluseviciute, Rafael Riudavets-Puig, Romain Blanc-Mathieu, et al. Jaspar 2024: 20th anniversary of the open-access database of transcription factor binding profiles. *Nucleic Acids Res*, 52(D1):D174–D182, 2024.

Ramprasaath R Selvaraju, Michael Cogswell, Abhishek Das, et al. Grad-CAM: Visual explanations from deep networks via gradient-based localization. In *Proc IEEE Int Conf Comput Vis*, pages 618–626, 2017.

Leslie N Smith and Nicholay Topin. Super-convergence: Very fast training of neural networks using large learning rates. In *Artificial intelligence and machine learning for multi-domain operations applications*, volume 11006, pages 369–386. SPIE, 2019.
